# Complementing 16S rRNA gene amplicon sequencing with estimates of total bacterial load to infer absolute species concentrations in the vaginal microbiome

**DOI:** 10.1101/598771

**Authors:** Florencia Tettamanti Boshier, Sujatha Srinivasan, Anthony Lopez, Noah G. Hoffman, Sean Proll, David N. Fredricks, Joshua T. Schiffer

## Abstract

Whereas 16S rRNA gene amplicon sequencing quantifies relative abundances of bacterial taxa, variation in total bacterial load between samples restricts its ability to reflect absolute concentration of individual species. Quantitative PCR (qPCR) can quantify individual species, but it is not practical to develop a suite of qPCR assays for every bacterium present in a diverse sample. We analyzed 1320 samples from 20 women with a history of frequent bacterial vaginosis, who self-collected vaginal swabs daily over 60 days. We inferred bacterial concentrations by taking the product of species relative abundance (assessed by 16S rRNA gene amplicon sequencing) and total bacterial load (measured by broad-range 16S rRNA gene qPCR). Log_10_-converted inferred concentrations correlated with targeted qPCR (r = 0. 935, p<2.2e-16) for seven key bacterial species. The mean inferred concentration error varied across bacteria, with rarer bacterial vaginosis-associated bacteria associated with larger errors. 92% of errors >0.5 log_10_ occurred when relative abundance was <10%. Many errors occurred during early bacterial expansion or late contraction. When relative abundance of a species is >10%, inferred concentrations are reliable proxies for targeted qPCR. However, targeted qPCR is required to capture bacteria at low relative abundance, particularly with BV-associated bacteria during the early onset of bacterial vaginosis.

## Introduction

For most infectious diseases, the absolute concentration of a single pathogen is often the most specific marker of disease severity and therapeutic response(1–3). In constrast, studies of bacterial communities usually rely on broad-range consensus sequence PCR of taxonomically informative genes (such as 16S rRNA) coupled with next generation sequencing (NGS) to assess relative, but not absolute abundances of bacteria. At a mechanistic level, specific combinations of bacteria and bacterial gene products are thought to play a causative role in the pathogenesis of many microbiome associated conditions(4–6), and this approach of characterizing the microbiota is valuable. However, absolute concentration of individual bacterial taxa within communities may be a better predictor of biological activity or disease risk compared to relative abundances of these taxa. Quantitating absolute concentration of individual species with qPCR is time intensive, requires generation of a standard curve for each organism using known concentrations of DNA, is expensive and only available in specialized laboratories. Moreover, each qPCR assay requires significant development and validation costs. qPCR is therefore not typically comprehensive for all species in a community. Moreover, selection of the most appropriate species for analysis may reflect investigator bias.

A method to infer absolute concentration of multiple bacterial species from NGS data would be extremely useful for the field including studies of the vaginal microbiome. NGS amplicon sequencing is a fractional approach that has been used to help define conditions such as bacterial vaginosis (7–10), and to identify enhanced risk for other sexually transmitted infections and pre-term delivery (11,12). However, total bacterial load may vary significantly between and within individuals over time even over the course of a single day (8). Therefore, relative abundances may not accurately represent absolute concentrations. Consequently, as shown recently in the gut microbiome, relative abundances may identify spurious disease associations which may in fact be driven by total microbial load (13).

Here, we demonstrate that multiplying relative abundance data (composition) by estimates of total bacterial DNA as measured by a broad-range 16S rRNA gene qPCR assay provides useful estimates of absolute concentrations of bacterial DNA for a given species. These inferred concentrations have already been used in studies of the penile microbiome, though without formal validation (14). Herein we validate inferred concentrations by comparison of absolute concentrations measured by targeted qPCR assay for seven key species in the vaginal microbiome. We find that whereas inferred concentrations are accurate for most samples, they are prone to error when relative abundance is low and may misrepresent kinetics of individual species during critical periods of expansion and clearance.

## Matherials and Methods

### Ethics statement

Vaginal samples were collected using protocol 417, which was approved by the institutional review board (IRB) at the University of Washington (approval no.: STUDY00000398). All participants provided written informed consent prior to study enrollment. Consent forms were approved by the IRB as part of protocol 417.

### Study Population

The study population was comprised of 20 women enrolled in a longitudinal study of bacterial vaginosis (BV) natural history at the University of Washington Virology Research clinic between 2015 and 2017. At enrollment, participants were given sufficient swabs for three times daily swabs over 60 days. Diagnosis, sample collection, storage, and processing of swabs are as described in (15). Participants were also given a study diary to record symptoms of BV, antibiotic use, menstruation, sexual activity and other medical events. In total, as some participants occasionally skipped samples, we analyzed 1320 data points for each of the seven key species.

### DNA Extraction and Quantitative Polymerase Chain Reaction (qPCR)

DNA was extracted from vaginal swabs using the BiOstic Bacteremia DNA Isoaltion Kit (Mobio, Carlsbad, CA). Sham swab without human contact were extracted in parallel to assess contamination from reaction buffers or the collection swabs. No template water controls were included to determine if there was any contamination from PCR reagents. Each sample was evaluated for PCR inhibition (Khot et. al. BMC Infectious Diseases. 2008) and total bacterial concentrations in each sample were measured using a qPCR assay that targets the V3-V4 region of the 16S rRNA gene (Srinivasan et al. PloS ONE 2012). Concentrations of specific vaginal bacteria were measured using qPCR assays targeting 7 key vaginal bacteria: *Atopobium vaginae*, BV-associated bacterium 2 (BVAB2), *Gardnerella vaginalis, Lactobacillus crispatus, Lactobacillus jensenii, Lactobacillus iners*, and *Megasphaera* (combined species 1 and 2) species (12,16,17). We measured relative abundances of bacterial taxa using broad-range PCR targeting the V3-V4 region of the 16S rRNA gene with next-generation sequencing on the Illumina MiSeq instrument (Illumina, San Diego, CA)(18). The *DADA2* pipeline was used to infer sequene variants from raw reads for subsequent analysis (19). Sequences were classified using the phylogenetic placement tool *pplacer* (20) and a curated reference set of vaginal bacteria (8). In subsequent text, we use NGS to to refer to data generated using broad-range PCR and sequencing. Sequence reads have been submitted to the NCBI Short Read Archive (in submission. Accession numbers pending). Relative abundances and absolute concentrations of specific vaginal bacteria were measured on all samples in two participants and in daily morning samples for the remaining 18. We performed qPCR on all samples collected from each participant, but for the purpose of this work only consider the morning samples.

All data generated or analysed during this study are included in the supplementary material.

### Statistical considerations

We calculated inferred concentrations using equation 1.

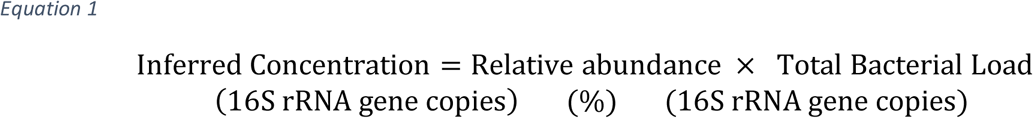

We present the plots and related calculations on a log_10_ scale. To keep all values finite, zero relative abundance (%) were mapped to 1e-5 and zero inferred concentrations were mapped to 1. The choice of this mapping changes some of the numerical results presented here, namely the correlation coefficient and the clustering class of the samples. However, the general observations being made are consistent.

We defined the error of inferred concentration, IC error, as

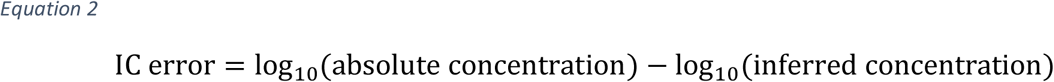

Rates of change per day where calculated between any two consecutive time points which were 18-36 hours apart. Rates were calculated from log_10_ converted values for relative abundance and inferred and absolute concentration. We defined the error in rates from inferred concentrations, rIC error, as

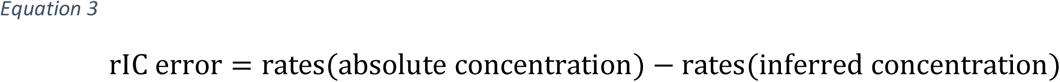

Comparison of means was done using the t.test function in R (21). We used Pearson’s correlation coefficient for all correlation analysis. This was done using the cor.test function in the stats package in R (21). Correlation coefficients were compared using the Cocor package in R (22). The suite provides 10 test for overlapping correlations, i.e. measurements taken from the same data set. All test were significant, but we report the value of the Hittner test here for simplicity. The Breusch-Pagan test was used to test the heteroskedasticity of the linear regression model of the relative abundance and inferred concentration vs absolute concentration. It tests whether the variance of the erros from a regression is dependent on the values of the independent variables. This was implemented using the bptest of the lmtest package in R (23).

We constructed the dendrograms for clustering analysis by complete linkage hierarchical clustering of species abundance and/or concentration based on Euclidean distance between all sample pairs.

We tested concordance between pairs of dendrograms using the entanglement coefficient found in the dendextend package in R (24). To calculate the coefficient, first all the samples are numbered in the order they appear for each tree. The coefficient is then calculated by taking the Euclidean distance of these two vectors which is then normalised by the worst case entanglement value (i.e. the Euclidean distance when the order of the two dendrograms is opposite). The entanglement coefficient thus defined ranges from 0 to 1, with 0 indicating perfect alignment between the dendorgrams and 1 a complete mismatch.

## Results

### Bacterial kinetics in 20 women with frequent recurrent BV

The bacterial kinetics observed for a single participant are shown in **Figure 1a and b.** The individual shown underwent dynamic changes in bacterial profile with notable shifts between low to high diversity states. The bacterial kinetics of 19 other participants can be found **in Figure S1**. As previously noted, high diversity states were often concurrent with high absolute concentrations of *Gardnerella vaginalis, Atopobium vaginae*, BVAB2 and *Megasphaera*, which all have been associated with bacterial vaginosis (BV) (8,10,25).

**Figure 1.**
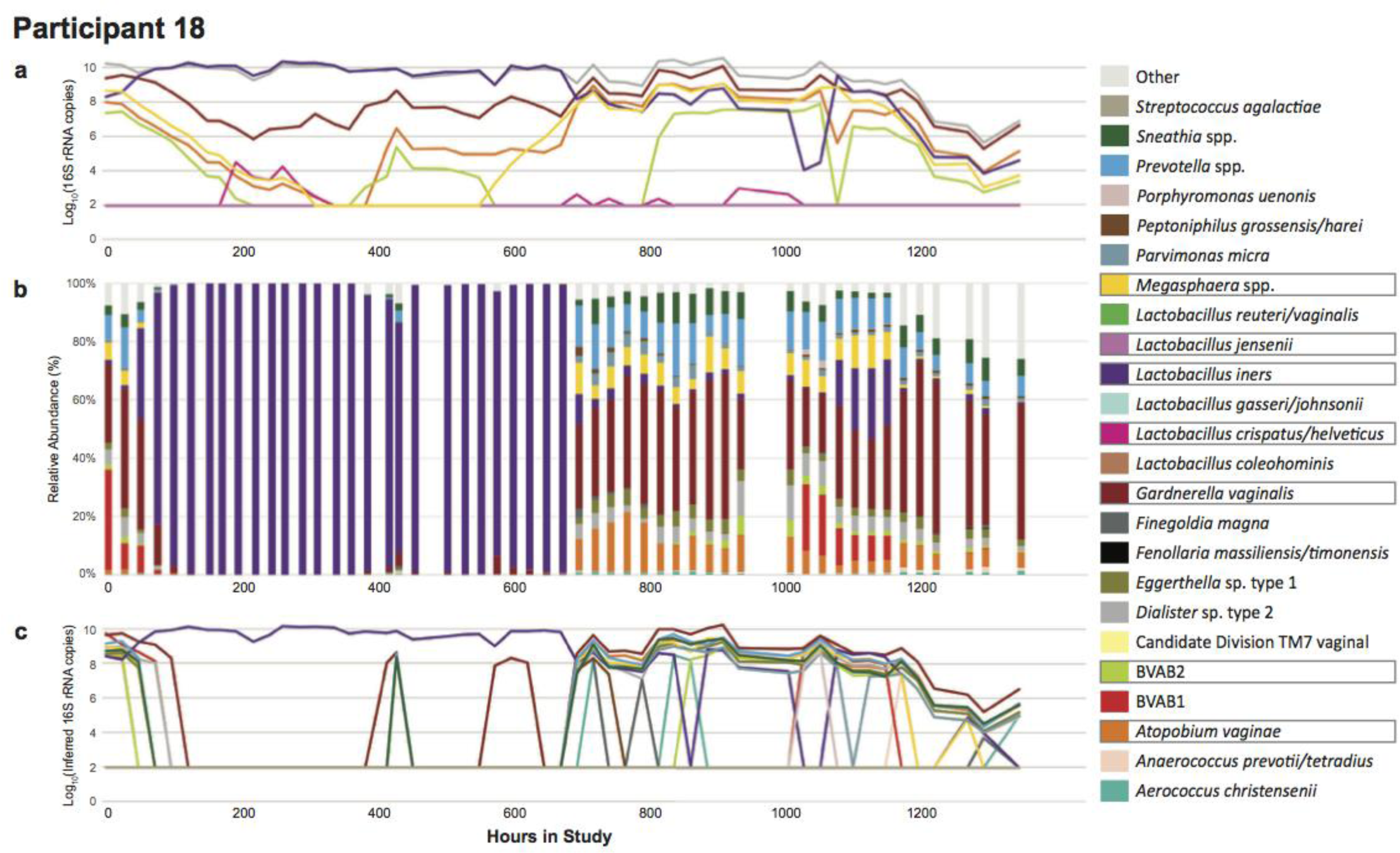
Complex bacterial kinetics in the vaginal niche in one representative study participant. Daily samples from a woman, Participant 18, who performed self-swabbing of the vagina were analyzed by: **a)** targeted qPCR of seven specific species, **b)** high throughput sequencing using 16S rRNA and **c)** inferred concentration for species with relative abundance above 1%. qPCR allows measures of absolute concentration, whereas broad range PCR with sequencing provides a measure of bacterial diversity in a given sample. Targeted qPCR often detects shifts in single species prior to NGS. Inferred concentration follows qPCR more closely than relative abundance does and may project concentration of species for which targeted qPCR assays are not available.

In 5 of the participants, shifts in composition appear less abruptly when measured by single species qPCR than by NGS. For example, for the participant shown in Figure 1, the absolute concentration of *A. vaginae* increases on day 17 (hour 415), but its relative abundance does not show a consistent increase until day 28 (hour 671) although there are some non-zero abundances in 4/9 samples before this point. From day 0 to day 28 (hour 168), the participant received metronidazole for BV: qPCR shows an exponential decline in BV-associated species absolute concentrations(26); yet, NGS shows a much more abrupt shift towards *L. iners* predominance. NGS can also fail to capture low-level colonization of bacteria, such as that of *G. vaginalis* on days 12 to 11 (hours 150 and 261). Several high-diversity samples have highly prevalent species which were not measured with qPCR in this study, such as *Prevotella bivia* from day 28 onwards (hour 671). These observations, which can be made for many of the individuals in this cohort, highlight that qPCR provides more granular estimates for measuring single species kinetics while NGS is optimal to estimate bacterial diversity in high diversity communities.

### Relative abundances may misclassify single species absolute concentration due to shifts in total bacterial load

We compared absolute concentration and relative abundance from the same samples measured within individuals over the course of the study. Examples for two species, *L. crispatus* and *Megasphaera*, are shown in **Figure 2a** and **b** (examples for the remaining five species are in supplementary **Figure S2**). There were time points in which absolute and relative abundance measures demonstrated opposing or differing kinetics, often due to concurrent large shifts in total bacterial load. These are indicated by arrows in **Figure 2a** and **b**. Thus, relative abundance may misrepresent absolute concentration when not accounting for total bacterial load.

**Figure 2.**
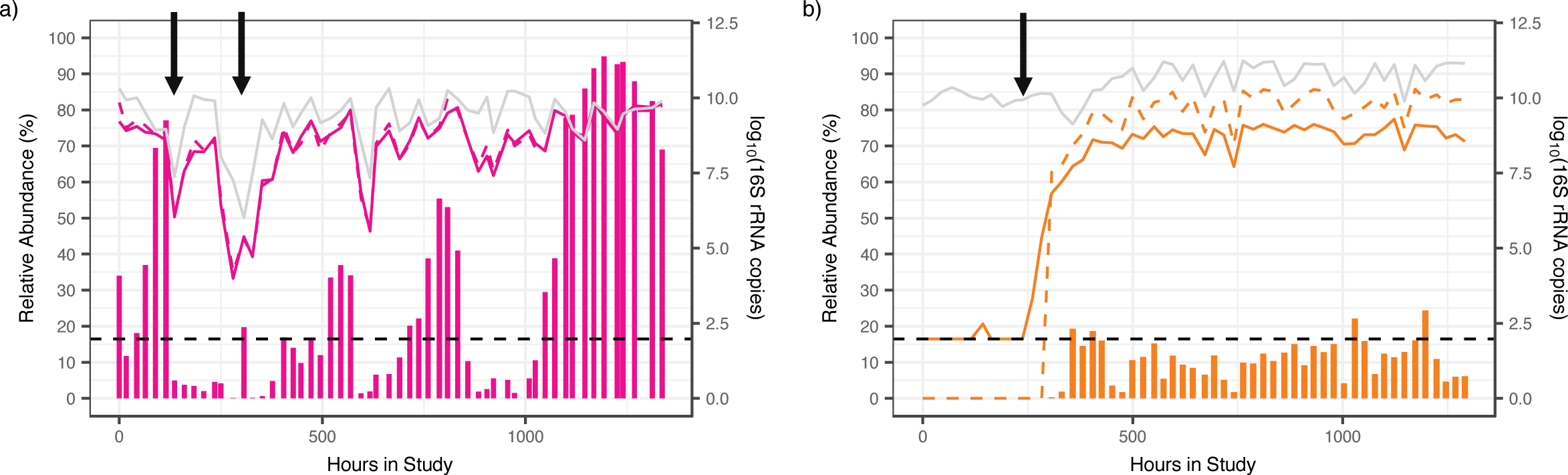
Relative abundances estimates can misrepresent actual concentrations due to shifts in total bacterial load. Examples of species-specific profiles in two participants for two different species **a)** *L. crispatus*, Participant 06 and **b)** *Megasphaera*, Participant 17. Vertical bars show relative abundance (%, left-y-axis), solid lines are absolute concentrations measured by qPCR and grey line is total bacterial load, dashed lines are inferred concentrations (all right y-axis). The dashed black line indicates detection threshold for qPCR data (93.8 16S rRNA copies). Arrows indicate timepoints when relative abundance changes are discordant from absolute concentration changes, which often occur when bacterial loads shift dramatically, or relative abundance is low. Examples for the remaining species can be found in **Fig S2.**

### Inferred concentrations are predictive of absolute concentrations measured by qPCR

For each species we calculated inferred concentrations by multiplying total bacterial load by NGS-relative abundance as shown in equation 1. We then compared these with absolute concentration as measured by targeted qPCR assay for the seven key species. For each species, inferred bacterial concentration closely tracked absolute concentration for most samples **(dotted line in Figure 2a** and **b and Figure S2)**. In many instances and for most species there were no obvious extreme discordance noted (**Figure 2a** and **S2**). For some species however, such as *Megasphaera* and BVAB2, inferred concentration consistently overestimated absolute concentration by an order of magnitude (**Figure 2b** and **S2d**). In a subset of samples, for all species, inferred concentration was zero while qPCR levels were positive leading to profound discordance between inferred and absolute concentration: this was most often noted at low absolute concentration **(Figure 2a and b)**.

We compared correlation between relative abundance and absolute concentration (r=0.936, P < 2.2e-16 **Figure 3a**) to correlation between inferred concentration and absolute concentration (r=0.935, p<2.2e-16 **Figure 3b**). The two correlation coefficients are not statistically different (Hittner test, p>0.08)(22). Species-specific correlations were noted. For inferred concentrations, *Megasphaera* and BVAB2 produced the strongest correlation followed by *L. crispatus, A. vaginae* and *L. jensenii*; *G. vaginalis* and *L. iners*, which are often present at moderate concentrations (~10^6^16S rRNA gene copies), had the weakest correlations though correlations coefficients for all species were high **(Table 1)**.

**Table 1.** Pearson correlation coefficients of single species between absolute concentration versus relative abundance and inferred concentration.

| Species | PCC Relative Abundance | PCC Inferred Abundance |
| --- | --- | --- |
| Megasphaera | 0.969 | 0.978 |
| BVAB2 | 0.942 | 0.952 |
| Lactobacillus crispatus | 0.941 | 0.920 |
| Atopobium vaginae | 0.911 | 0.916 |
| Lactobacillus jensenii | 0.908 | 0.911 |
| Gardnerella vaginalis | 0.881 | 0.890 |
| Lactobacillus iners | 0.863 | 0.872 |

**Figure 3.**
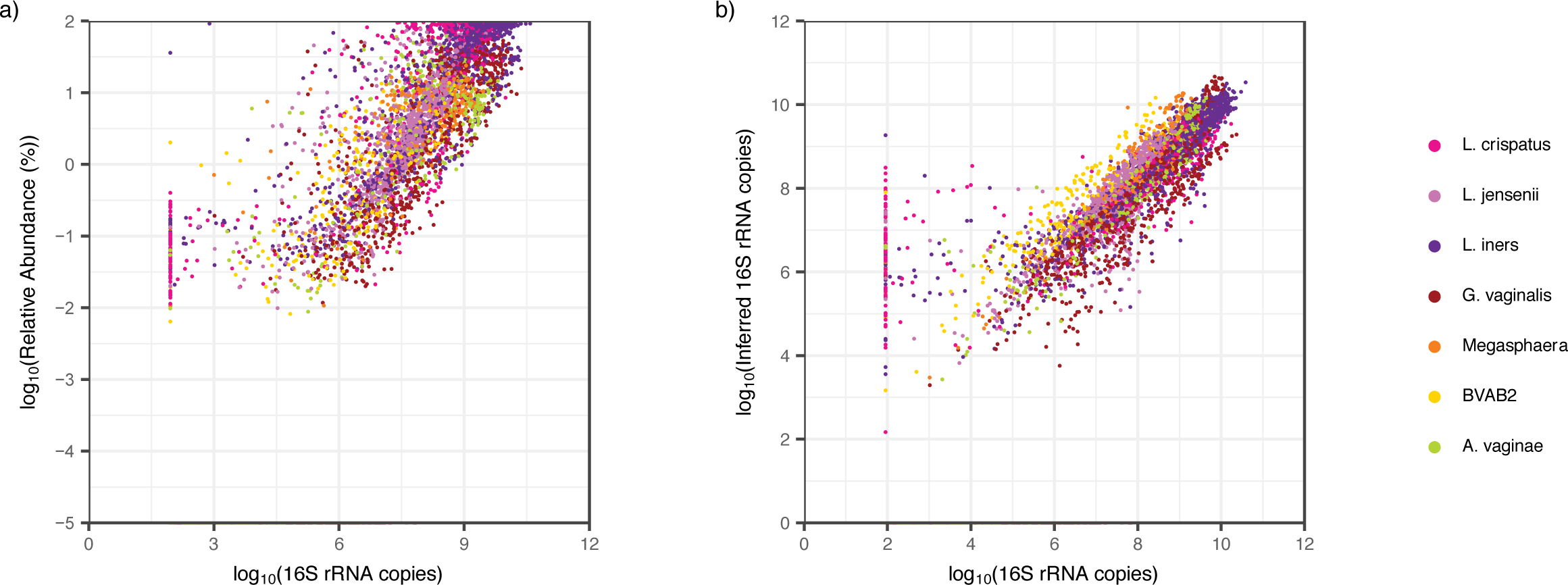
Inferred concentration correlates more strongly with absolute concentration than relative abundance. **a)** Scatter plot of relative abundance vs absolute concentration. Pearson correlation coefficient (pcc), r is 0.936, P<2.2e-16. **b)** Scatter plot of inferred concentration vs absolute concentration. Both axes are plotted on a logarithmic scale. Pearson correlation coefficient (pcc), r is 0.935, P<2.2e-16. Samples which were negative by NGS but not by targeted qPCR are plotted on the x-axis while samples negative by targeted qPCR but positive by NGS are listed on the reported threshold for targeted qPCR (log_10_(93.8) 16S rRNA gene copies). Relative abundances and inferred concentrations were generally falsely negative at low absolute concentrations. Variance in the relationship between absolute concentration and relative abundance is inversely proportional to species concentrations (Breusch-Pagan test, P=2e-3). Whereas this relationship was not statistically significant between absolute concentration and inferred abundance (Breusch-Pagan test, P=0.06).

We defined error of inferred concentration, IC error, as in equation 2. While there was a large range in errors for non-zero inferred concentrations (**Figure 3b**, range: −7.32 log_10_ (16S rRNA gene copies) – 2.66 log_10_(16S rRNA gene copies)), the mean IC error (−0.319 log_10_(16S rRNA gene copies)) and standard deviation (0.999 log_10_(16S rRNA gene copies)) were low. Moreover, the median IC error for most species approximated zero with samples within the interquartile range demonstrating minimal IC error (**Figure 4a**). However, for BVAB2 and *Megasphaera*, the interquartile range of IC error, while narrow, was all less than zero, implying consistent overestimation of absolute concentration by inferred concentration (pair-wise t-test p<0.05). There was a trend towards global underestimation of *G. vaginalis* using inferred concentration **(Figure 4a)**.

**Figure 4.**
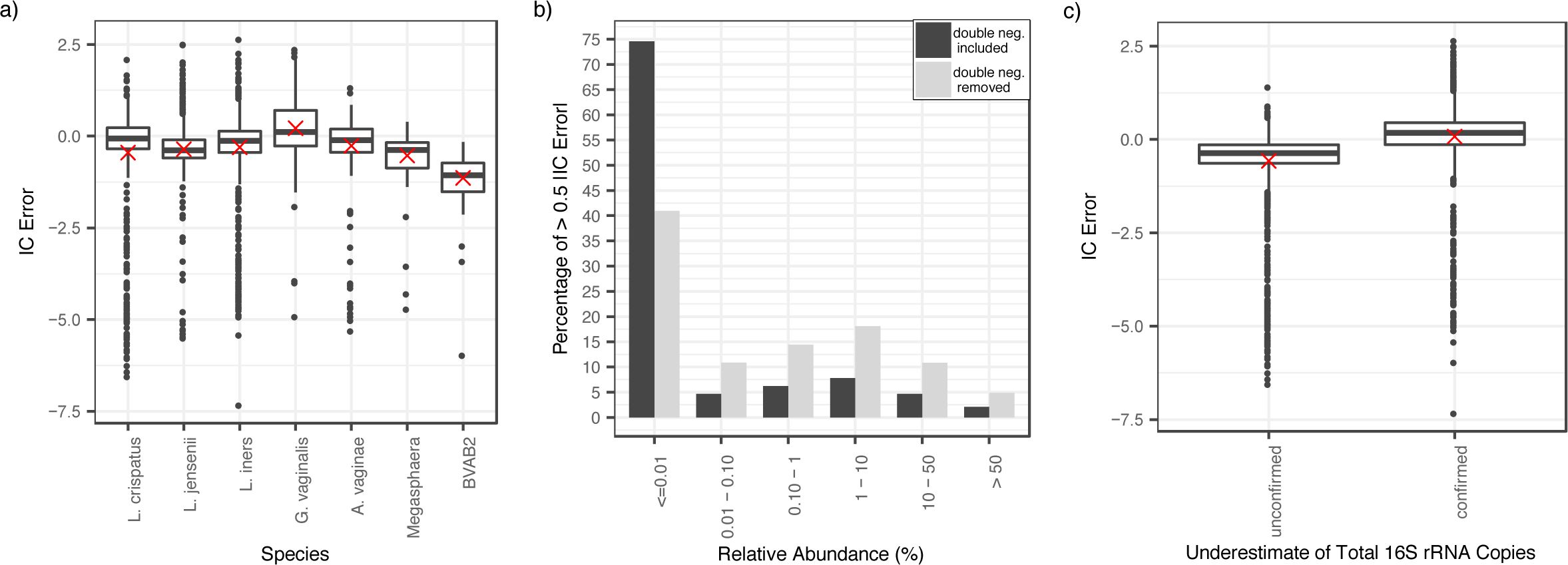
Low relative bacterial abundance is the major predictor of IC error for inferred concentrations compared to absolute concentrations. **a)** Boxplots displaying IC error (equation 2), with zero inferred concentrations removed, indicate low IC error rates overall. Inferred values are consistent overestimates for BVAB2 and *Megasphaera* spp. Boxes are the interquartile range; whiskers are 1.5x the IQR and dots are samples outside of this range; red crosses are mean. **b)** Bar-chart of incidence of >0.5 IC error by relative abundance group. Black is inclusive of double-negatives (0 inferred concentration and threshold absolute concentration): 93% of >0.5 IC errors are accounted for by relative abundances <10% (85% by relative abundances <1%). In grey, concurrent negative samples are removed: 84% of >0.5 IC errors are accounted for by relative abundances <10% (66% by relative abundances <1%). **c)** Boxplots displaying IC error for samples with unconfirmed and confirmed underestimates of total bacterial load by broad range qPCR assay (samples where BR16S is lower than the sum of concentrations of the seven targeted species). Data points with zero inferred concentration were removed. Samples with underestimates of total bacterial load overestimate concentration more than other samples. Overall however, the range of IC error is comparable between both groups. Boxes are the interquartile range; whiskers are 1.5x the IQR and dots are samples outside of this range; crosses are mean.

### Low relative abundance is the major source of IC error

The variance in the relationship with absolute concentration tended to be inversely proportional to species concentrations [Breusch-Pagan test; p = 0.06] highlighting that a larger range of IC errors tended to be reported at lower bacterial loads (**Figure 3a**). Accordingly, 93% of >0.5 IC errors were accounted for by relative abundances below 10% and 85% by relative abundances below 1%. Many of these IC errors occurred on double negatives – samples for which inferred concentration was zero and absolute concentration was reported at threshold. When these samples were removed from the analysis, 84% of >0.5 IC errors were accounted for by relative abundances <10% and 66% by relative abundances below 1% **(Figure 4b).** The median absolute concentration above the limit of detection for >0.5 IC errors was 5.95 log_10_(16S rRNA gene copies) (IQR: 4.03 – 7.88, range: 1.97 – 10.39).

We defined false positive samples as non-zero inferred concentration values when absolute concentration qPCR values were at or below the detection threshold and false negatives as zero-values for inferred concentration when absolute concentrations were above the detection threshold. False negatives were more common (23.6% of samples) than false positives (3.17 % of samples) which demonstrates that targeted qPCR is more sensitive for single species detection than NGS.

The incidence of false negatives was not equal across species, with *G. vaginalis* having the highest percentage of false negatives, followed by *L. inners* and *A. vaginae* (*L*.*crispatus* 13.8%, *L. jensenii* 31.1 %, *L. iners* 35.1%, *G. vaginalis* 60.4%, *A. vaginae* 35.3%, *Megasphaera* 5.40%, BVAB2 9.84%). The higher percentages of false negative for some species occurred because they are often present at moderate concentrations, near the relative abundance error threshold. The median qPCR value for false negative samples was 3.92 log_10_ (16S rRNA gene copies) (IQR: 2.88 – 4.82, range: 1.97 −7.84), again showing that IC errors generally occur at lower bacterial loads.

Total bacterial load measured by broad-range qPCR assay was frequently below the sum of the concentration of all seven species measured by targeted qPCR assays (37.6% per species per sample). Non-zero inferred concentrations from samples with underestimates of total bacterial load consistently overpredicted absolute concentration (one-tailed t-test p<2.6e-4) and did so more than at other points (pair-wised t-test p<2.2e-16) (**Figure 4c**). Non-zero inferred concentrations from samples with known underestimates of total bacterial load had a median IC error of 0.171 log_10_ (16S rRNA gene copies) (IQR −0.138 − 0.447, range −7.31 – 2.66) compared to −0.368 log_10_ (16S rRNA gene copies) (IQR −0.638 - −0.143, range: −6.54 – 1.42) in other samples.

*L. crispatus* had the highest percentage of false positives (*L*.*crispatus* 8.42%, *L. jensenii* 1.08%, *L. iners* 3.56%, *G. vaginalis* 0.46%, *A. vaginae* 3.07%, *Megasphaera* 1.12%, BVAB2 1.79%). The median relative abundance of false positives across all samples was extremely low 0.06% (IQR: 0.04 – 0.11%, range 0.0007 – 36.8%).

### Concentrations inferred from NGS predicts observed absolute concentration regardless of sample diversity or sequencing depth

Inferred concentrations did not disproportionally record misleading results from low or high diversity samples as measured by Shannon Diversity index (**Figure 5a**). Moreover, we observed occasional large absolute IC errors across all sequencing depths (**Figure 5b**). Low bacterial abundance was the primary source of absolute IC error regardless of diversity or sequencing depth **(Figure 5a** and **b)**. Larger than 0.5 absolute IC error was observed across all raw species counts, but the largest absolute IC errors (above >2) were almost exclusively associated with raw species counts below 100 **(Figure 5c)**.

**Figure 5.**
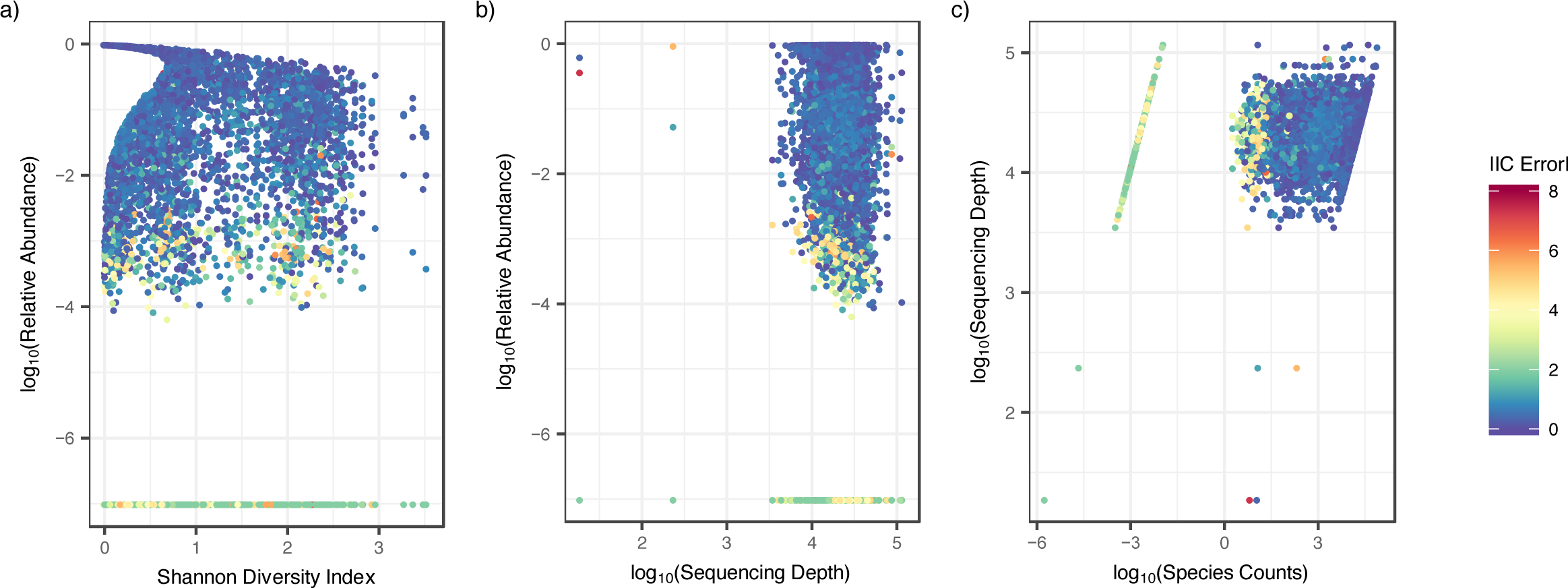
Sample diversity, sequencing depth and species counts do not impact IC error of inferred concentration. Scatter-plots color-coded by IC error. Each dot is a sample for a specific species from a single participant. **a)** Relative abundance versus Shannon diversity index. High IC error predominately occurred at low relative abundance but across both low and high diversity samples. **b)** Relative abundance versus sequencing depth. High IC error predominately occurred at low relative abundance but across various levels of sequencing depth. c) Sequencing depth vs species counts. High IC error occurred at species counts below 100, although >0.5 IC error is observed across all species counts.

### Inferred concentration estimates are predictive of most temporal changes in single species bacterial load

We examined whether inferred concentration is a useful tool for evaluating individual species kinetics by determining changes in bacterial levels over the course of a day. Rates of change in relative abundances correlated only weakly with absolute concentrations [r=0.271, p<2.2e-16]. Moreover, 17.1% of the time, we observed a change in relative abundance in the opposite direction to that of absolute concentration (top-left and bottom-right quadrants in **Figure 6a**). This type of error occurred commonly for both the most abundant (e.g. *L. crispatus*) and rarer species (e.g. BVAB2).

**Figure 6.**
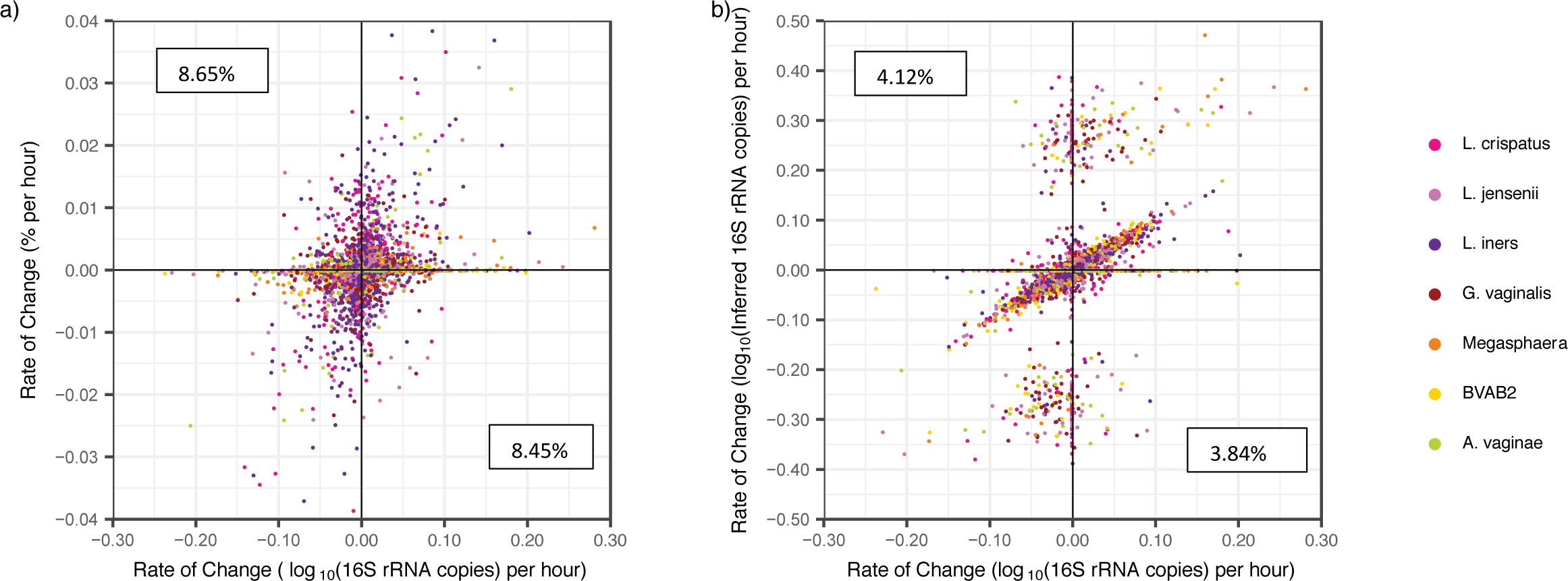
Inferred concentrations allow somewhat accurate inference of kinetic changes between two sequential samples. **a)** Scatter plot of change in relative abundance vs change absolute concentration shows poor correlation. Pearson correlation coefficient (pcc), r is 0.271 (P<2.2e-16). A high percentage of observed changes in relative abundance are in the opposite direction as those in absolute concentration (left upper and right lower error quadrants marked with percentages) **b)** Scatter plot of inferred concentration vs absolute concentration shows improved correlation. Both axes are plotted on a logarithmic scale. Pearson correlation coefficient (pcc), r is 0.392 (P<2.2e-16). Percentages correspond to the number of data points which fall within the error quadrants and are lower than for relative abundance. Inferred values misreport direction of kinetics less frequently.

Rates of change in inferred concentration showed improved correlation with rates of change in absolute concentration [pmcc=0.392, p<2.2e-16]. The mean rIC error (defined in the **Methods**) was low (−2.71 x10^-3^, SD: 1.54 log_10_(16S rRNA gene copies) per hour) though the range of rIC errors was high (−9.29 – 9.31 log_10_(16S rRNA gene copies) per hour), indicating occasional samples with very poor prediction. Inferred concentrations decreased the sign rIC error rate by more than 50% (from 17.1% to 7.97%, **Figure 6b**).

**Figure 7a** shows a typical profile of *A. vaginae* absolute levels and sample-to-sample change, to demonstrate the two types of rIC errors which were most common to the data. The first were large positive or negative rates which occurred when one of two consecutive points had an inferred concentration of zero, while absolute concentration was detectable by qPCR. These points resulted in dramatic overestimation of growth or contraction rates for individual species across all samples (**Figure 6b & 7b**, right upper and left lower quadrants). Such rIC error often occurred when species were transitioning to or from a low abundance (<10^6^16S rRNA gene copies per sample). The second type of rIC error, occurred when two consecutive points had inferred concentrations of zero, resulting in underestimation of growth or contraction rates for individual species **(Figure 7b)**. This phenomenon also commonly occurred when a species was transitioning to or from a low abundance (<10^6^16S rRNA gene copies per sample). These two forms of transitions accounted for 91.7% of rIC error > 0.05 **(Figure 7c)**. If all transitions involving a zero value were eliminated from the analysis, we observed excellent correlation between inferred and observed rate of change (r=0.876, p<2.2e-16). It follows that inferred concentrations do not capture kinetics during microbial blooming or contraction when bacteria are at low concentration or not detected using the less sensitive broad-range PCR with NGS approach. However, inferred concentrations can be used to estimate individual species growth and contraction rates when bacteria are present at higher concentrations such as >10^6^16S rRNA gene copies/swab.

**Figure 7.**
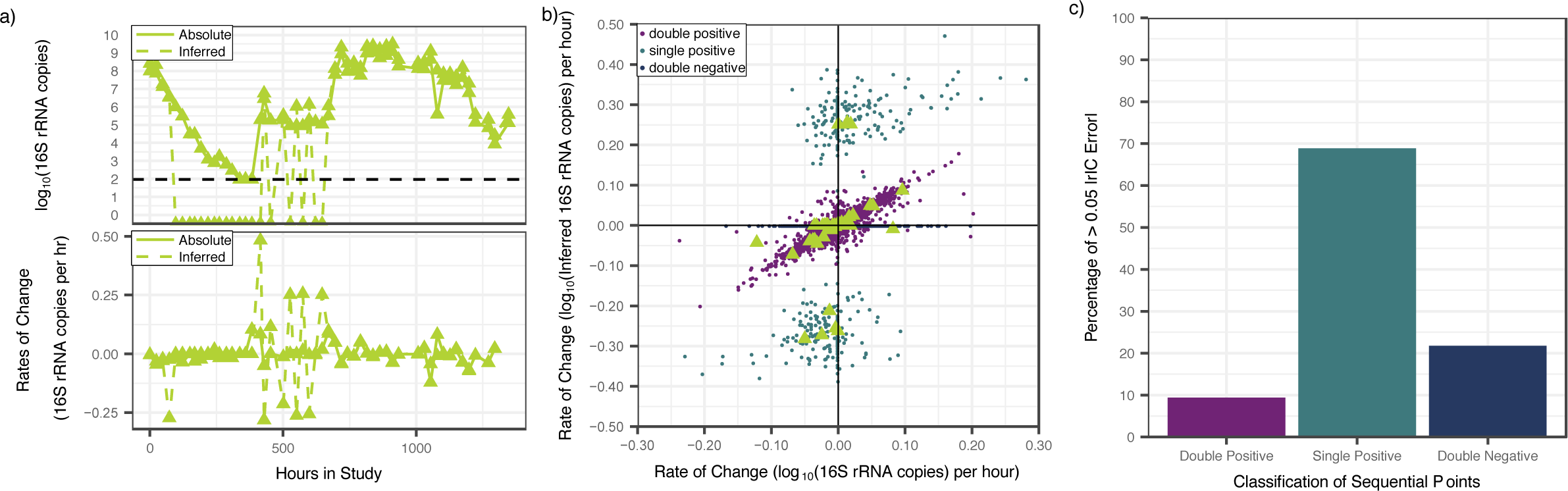
Inferred concentration measures allow accurate inference of kinetic changes between two sequential non-negative samples. **a)** Top: Levels of *A. vaginae* over time in a single participant (dotted is inferred and solid is absolute concentration); bottom: rate of change in levels of *A. vaginae* over time in the same participant (dotted is inferred and solid absolute concentration); divergence in swab to swab levels between inferred and absolute concentrations varies only when inferred concentration is zero in one of the sequential samples. **b)** Scatter plot of rate of change of inferred concentration as predicted by NGS vs qPCR observed values. Both axes are plotted on a logarithmic scale. Data is the same as in **Fig 6 a** and **b**. Triangles correspond to panel **c**. Points are colored according to whether consecutive samples were double positive (both >0 inferred concentration), single positive (one >0 and one 0 inferred concentration) or double negative (both 0 inferred concentration). Data points in which both samples are positive (no zeroes) are much more highly correlated (r is 0.876, P<2.2e-16). **c)** A majority of rIC errors > 0.05 occur during transitions between positive and negative samples (one zero).

### Complete linkage clustering by inferred and absolute concentrations shows general agreement

To assess whether inferred concentrations provide similar or disparate classification of samples, we clustered samples using complete linkage hierarchical clustering based on Euclidean distances (21) by inferred and absolute concentrations **(Figure S3)**. We compared the resulting dendrograms using the entanglement coefficient from the dendextend package in R (24), where a value of 1 corresponds to complete discordance and a value of 0 indicates perfect alignment. The two dendrograms were found to be in agreement, with a low entanglement coefficient 0.11.

We next determined the number of clusters using NbClust package in R (27). Absolute concentration identifies two whereas inferred concentration identifies three clusters. The third cluster arose from a general disctinction between samples dominated by *L. crispatus* from *L. iners* as the inferred concentrations had a lower threshold (1 copy per swab) than the qPCR (93.8 copies per swab).

### Inferred concentration may provide the most comprehensive overview of individual species kinetics

Inferred concentrations can be calculated for all species captured by NGS. In **Figure 1** and **S1**, we show the inferred concentrations of the most abundant species across all samples. We imposed a 1% relative abundance threshold to limit the possible 0.5 IC error described in **Figure 4b**. This relative abundance cut-off results in abrupt appearance and disapparence of organisms. Although we cannot validate our projections for species outside of the seven key bacterial species for which we have targeted qPCR assays, inferred concentrations have the potential to describe the kinetics of relevant species present at moderate to high concentrations during bacterial shifts in the microbiome.

We carried out complete linkage hierarchical clustering based on Euclidean distance by inferred concentration and relative abundance for the 20 most abundant species of the data set **(Figure S4)**. The resulting dendrograms showed general agreement, with an entanglement coefficient of 0.12. Both techniques identified two clusters defined by high concentration *G. vaginalis* and high diversity versus *Lactobacillus* predominance (27).

## Discussion

An ideal assay that characterizes bacterial communities in an ecological niche would capture several metrics including species composition, diversity, and quantity as reflected by absolute concentration of all species present. Broad-range PCR of phylogenetically informative genes followed by NGS is the most commonly used approach and captures the first two metrics. However, because total bacterial levels may shift dramatically over narrow time intervals, relative abundance measures by NGS do not reflect absolute concentration. While it is possible to circumvent this issue with targeted (taxon specific) qPCR, these assays are expensive, time consuming and only available in specialized laboratories. Invariably, the absolute concentration of many relevant species is left unmeasured due to these constraints.

This measurement gap is of high relevance to clinical studies of the human microbiome in which total bacterial load may not be stable. It is biologically plausible that the absolute levels of critical species are more predictive of health and disease states than relative levels, as is the case with classical single pathogen infectious diseases. Moreover, serial measurements of absolute levels are necessary to fully capture non-linear microbial dynamic changes which relate to inter-species competition for limited resources.

Using a large longitudinal dataset of the vaginal microbiome notable for frequent changes between low and high diversity states, we demonstrate that the absolute concentration of a given species can be inferred by multiplying the total bacterial quantity by its relative abundance as measured by NGS. Given that quantitating total bacterial load is affordable and available to many laboratories, this simple approach may allow estimation of absolute concentration without needing to perform qPCR on all samples.

Our technique is remarkably predictive of absolute concentration with certain key exceptions. Species such as BVAB2 and *Megasphaera* which were often present at low absolute concentration were notable for high precision but slight inaccuracy: inferred concentration consistently slightly overestimated the abundance for these species. This result highlights that individual comparisons between inferred and absolute concentration must be performed for all species of interest. Other than in an exploratory fashion, we do not advocate the use of inferred concentration for species which have not been validated in depth with targeted qPCR assays and compared to absolute concentration.

Second, our approach has a very high IC error rate when relative abundance is low, or zero. In our qPCR dataset low level colonization of certain species often precedes a surge in levels prior to this species predominating. Because qPCR is more sensitive than NGS for low amounts of bacterial DNA, and because inferred concentration relies on NGS, inferred concentration will often miss persistent low-level colonization, as well as the critical early growth phase or late contraction phase of relevant species. Despite this fact, inferred concentration performs remarkably well at estimating growth and decay rates at the single species level, provided these rates are estimated based on positive sequential samples. One might be able to improve accuracy of the inferred concentrations by increasing sequencing depth or improving the accuracy of measurements of the total bacterial load.

A final issue not addressed by our technique is the limitation inherent to comparing bacterial quantities between species using qPCR based on differing amplification efficiencies of different assays. This variability may arise from different bacterial targets having varying GC content, secondary structures and amplification product size. In this sense, absolute concentration by qPCR may not be a perfect gold standard for comparing inferred concentration.

In summary, we developed and validated a simple, user-friendly method to estimate absolute species abundance in complex polymicrobial communities. This method is best employed when species are present at >10% relative abundance and must be validated for each species of interest. Ultimately, inferred concentration of one or several species may serve as a more predictive variable of disease association, compared to relative abundance, and may advance our understanding of how specific environmental and host factors influence microbial concentrations.

## Acknowledgements

This work was supported by the Sexually Transmitted Infections Cooperative Research Centers program (grant U19 AI 113173).

## Author Contributions

J.T.S, S.S. and D.N.F conceived and designed the experiments. A.L. performed the experiments. N.G.H. managed the NGS bioinformatic pipeline. S.P. managed data integration and contributed to figure generation. J.T.S. and F.A.T.B. conceived the idea of inferred concentration. F.A.T.B completed the analysis, contributed to figure generation and wrote the manuscript.

## Competing Interests

The authors declare no competing interests.

## References

1. Falade-Nwulia OO, Naggie S, Nahass RG, Kim AY, Scott JD, Ghany MG, et al. Hepatitis C Guidance 2018 Update: AASLD-IDSA Recommendations for Testing, Managing, and Treating Hepatitis C Virus Infection. Clin Infect Dis. 2018;67(10):1477–92.

2. File TM. Highlights from international clinical practice guidelines for the treatment of acute uncomplicated cystitis and pyelonephritis in women: A 2010 update by the infectious diseases society of America and the european society for microbiology and infectious. Infect Dis Clin Pract. 2011;19(4):282–3.

3. Saag MS, Benson CA, Gandhi RT, Hoy JF, Landovitz RJ, Mugavero MJ, et al. Antiretroviral drugs for treatment and prevention of HIV infection in adults: 2018 recommendations of the international antiviral society-USA panel. JAMA - J Am Med Assoc. 2018;320(4):379– 96.

4. Bisgaard H, Hermansen MN, Buchvald F, Loland L, Halkjaer LB, Bonnelykke K, et al. of the Airway in Neonates. N Engl J Med. 2007;357:1487–95.

5. Dejea CM, Fathi P, Craig JM, Boleij A, Taddese R, Geis AL, et al. Patients with familial adenomatous polyposis harbor colonic biofilms containing tumorigenic bacteria. Science (80-). 2018;359(6375):592–7.

6. Costello SP, Hughes PA, Waters O, Bryant R V., Vincent AD, Blatchford P, et al. Effect of Fecal Microbiota Transplantation on 8-Week Remission in Patients with Ulcerative Colitis: A Randomized Clinical Trial. JAMA - J Am Med Assoc. 2019;321(2):156–64.

7. Srinivasan S, Morgan MT, Fiedler TL, Djukovic D, Hoffman NG, Raftery D, et al. Metabolic signatures of bacterial vaginosis. MBio. 2015;6(2):1–16.

8. Srinivasan S, Hoffman NG, Morgan MT, Matsen FA, Fiedler TL, Hall RW, et al. Bacterial communities in women with bacterial vaginosis: High resolution phylogenetic analyses reveal relationships of microbiota to clinical criteria. PLoS One. 2012;7(6).

9. Ravel J, Gajer P, Abdo Z, Schneider GM, Koenig SSK, McCulle SL, et al. Vaginal microbiome of reproductive-age women. Proc Natl Acad Sci [Internet]. 2011;108(Supplement_1):4680–7. Available from: http://www.pnas.org/cgi/doi/10.1073/pnas.1002611107

10. Gajer P, Brotman RM, Bai G, Sakamoto J, Schütte UME, Zhong X, et al. Temporal dynamics of the human vaginal microbiota. Sci Transl Med [Internet]. 2012;4(132):132ra52. Available from: http://stm.sciencemag.org/content/scitransmed/4/132/132ra52.full

11. Nelson DB, Hanlon A, Nachamkin I, Haggerty C, Mastrogiannis DS, Liu C, et al. Early pregnancy changes in bacterial vaginosis-associated bacteria and preterm delivery. Paediatr Perinat Epidemiol. 2014;28(2):88–96.

12. McClelland RS, Lingappa JR, Srinivasan S, Kinuthia J, John-Stewart GC, Jaoko W, et al. Evaluation of the association between the concentrations of key vaginal bacteria and the increased risk of HIV acquisition in African women from five cohorts: a nested case-control study. Lancet Infect Dis [Internet]. 2018;18(5):554–64. Available from: http://dx.doi.org/10.1016/S1473-3099(18)30058-6

13. Vandeputte D, Kathagen G, D’Hoe K, Vieira-Silva S, Valles-Colomer M, Sabino J, et al. Quantitative microbiome profiling links gut community variation to microbial load. Nature [Internet]. 2017;551(7681):507–11. Available from: http://dx.doi.org/10.1038/nature24460

14. Liu CM, Prodger JL, Tobian AAR, Abraham AG, Price LB. crossm Penile Anaerobic Dysbiosis as a Risk Factor for HIV Infection.: 1–10.

15. Mayer BT, Matrajt L, Casper C, Krantz EM, Corey L, Wald A, et al. Dynamics of persistent oral cytomegalovirus shedding during primary infection in ugandan infants. J Infect Dis. 2016;214(11):1735–43.

16. Fredricks DN, Fiedler TL, Thomas KK, Mitchell CM, Marrazzo JM. Changes in vaginal bacterial concentrations with intravaginal metronidazole therapy for bacterial vaginosis as assessed by quantitative PCR. J Clin Microbiol. 2009;47(3):721–6.

17. Srinivasan S, Liu C, Mitchell CM, Fiedler TL, Thomas KK, Agnew KJ, et al. Temporal variability of human vaginal bacteria and relationship with bacterial vaginosis. PLoS One. 2010;5(4).

18. Garcia K, Celustka K, Srinivasan S, Loeffelholz T, Fiedler TL, Aker S, et al. Stool Microbiota at Neutrophil Recovery Is Predictive for Severe Acute Graft vs Host Disease After Hematopoietic Cell Transplantation. Clin Infect Dis. 2017;65(12):1984–91.

19. Callahan BJ, McMurdie PJ, Rosen MJ, Han AW, Johnson AJA, Holmes SP. DADA2: High-resolution sample inference from Illumina amplicon data. Nat Methods. 2016;13(7):581– 3.

20. FA M, RB K, EV A. pplacer: linear time maximum-likelihood and Bayesian phylogenetic placement of sequences onto a fixed reference tree. BMC Bioinformatics [Internet]. 2010;11:538. Available from: http://dx.doi.org/10.1186/1471-2105-11-538

21. Team RC. R: A Language and Environment for Statistical Computing. Vienna, Austria: R Foundation for Statistical Computing; 2018.

22. Diedenhofen B, Musch J. Cocor: A comprehensive solution for the statistical comparison of correlations. PLoS One [Internet]. 2015;10(4):1–12. Available from: http://dx.doi.org/10.1371/journal.pone.0121945

23. Achim Zeileis TH. Diagnostic Checking in Regression Relationships. R News [Internet]. 2010;2(3):7–10. Available from: http://cran.r-project.org/doc/Rnews/

24. Sieger T, Hurley CB, Fiser K, Beleites C. Interactive Dendrograms: The *R* Packages **idendro** and **idendr0**. J Stat Softw [Internet]. 2017;76(10). Available from: http://www.jstatsoft.org/v76/i10/

25. Fredricks DN, Fiedler TL, Marrazzo JM. Molecular Identification of Bacteria Associated with Bacterial Vaginosis. N Engl J Med. 2005;353(18):1899–911.

26. Marrazzo JM, Fiedler TL, Srinivasan S, Mayer BT, Schiffer JT, Fredricks DN. Rapid and Profound Shifts in the Vaginal Microbiota Following Antibiotic Treatment for Bacterial Vaginosis. J Infect Dis. 2015;212(5):793–802.

27. Charrad M, Ghazzali N, Boiteau V, Niknafs A. Package ‘NbClust.’ 2015;9. Available from: https://cran.r-project.org/web/packages/NbClust/NbClust.pdf

