## Supplementary Figures for "Complementing 16S rRNA gene amplicon sequencing with estimates of total bacterial load to infer absolute species concentrations in the vaginal microbiome"

**Figure S1. Vaginal microbiome profile for all study participants.** Participants 08 and 09 show 8 hourly samples, remaining subjects show daily morning samples. **a)** targeted qPCR of seven specific species; **b)** high throughput sequencing using 16S rRNA for top 24 species across entire data set; **c)** inferred concentration for species with relative abundance above 1%. qPCR allows measures of absolute concentration, whereas broad range PCR with sequencing provides a measure of bacterial diversity in a given sample. Targeted qPCR often detects shifts in single species prior to NGS. Inferred concentration follows qPCR more closely than relative abundance does and may project concentration of species for which targeted qPCR assays are not available.

#### Participant 01

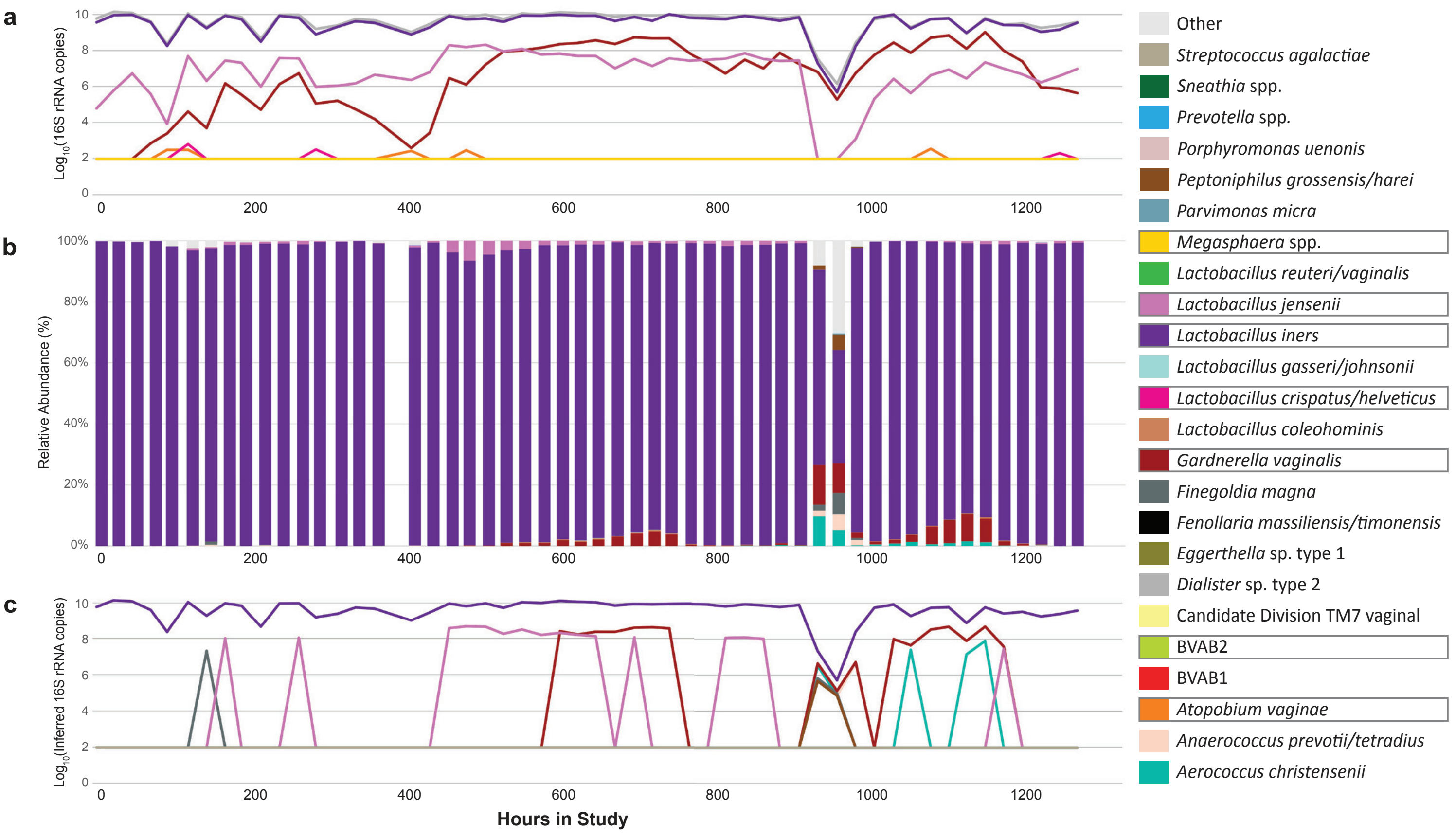


#### Participant 02

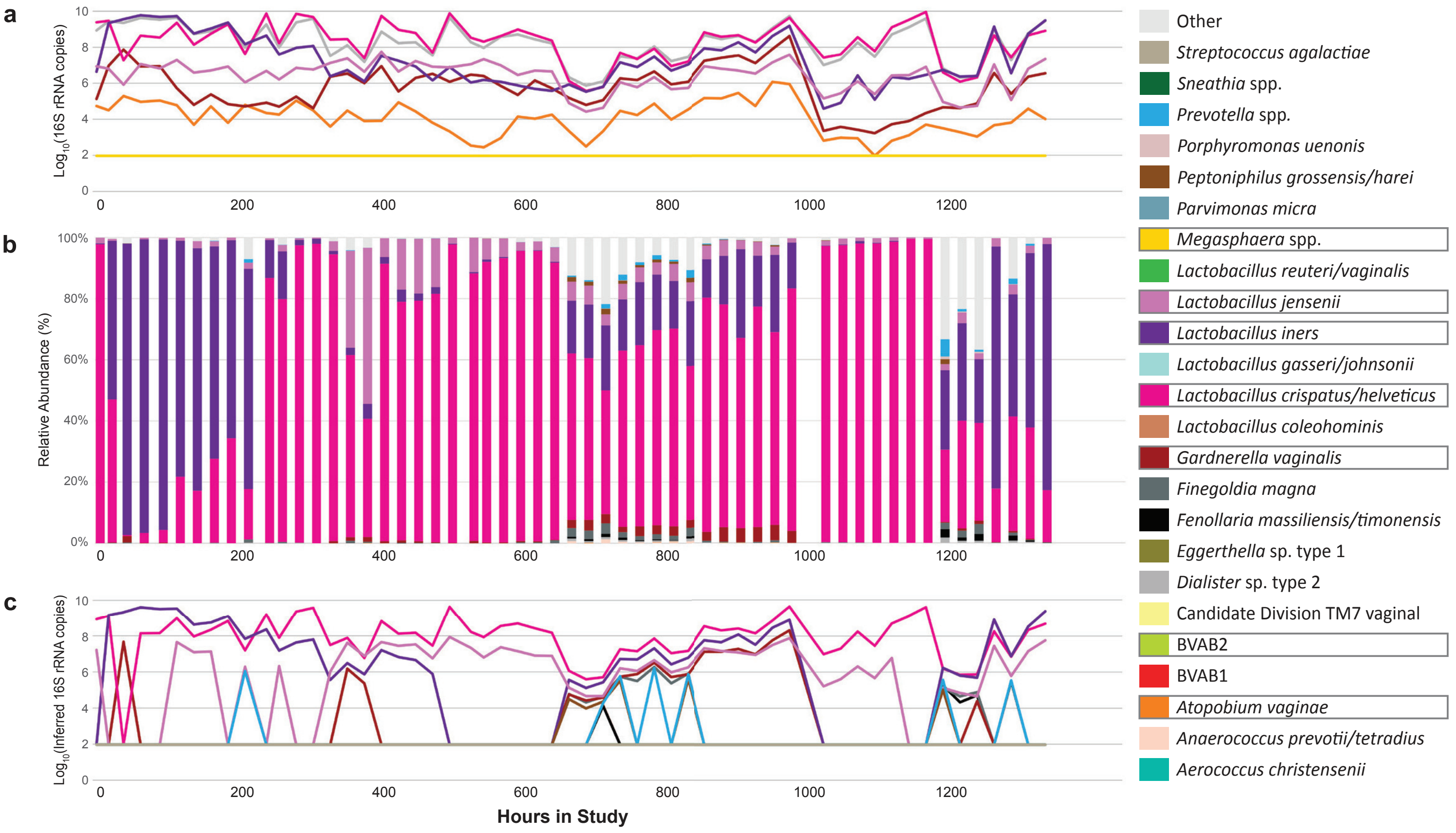

#### Participant 03

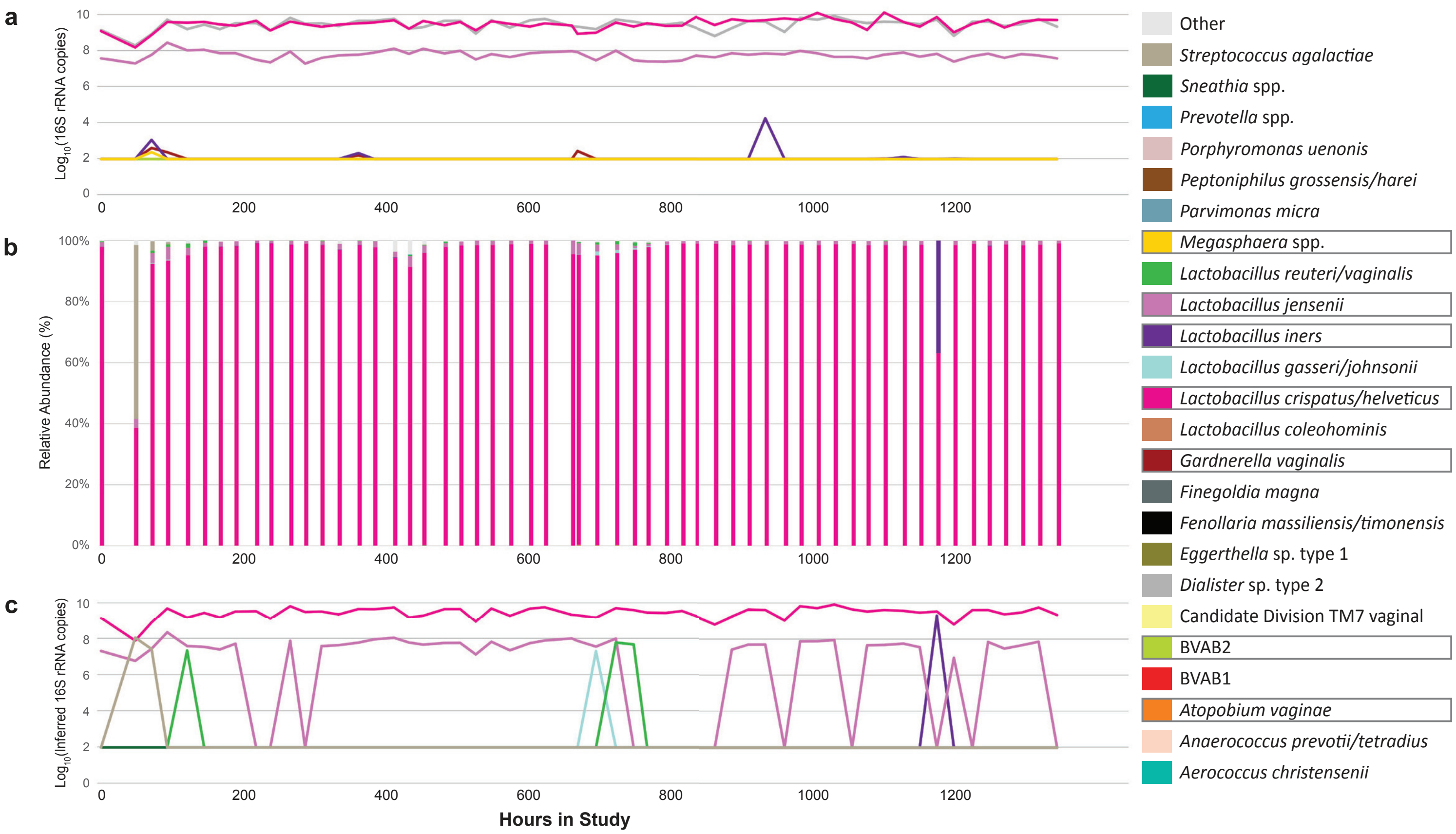


#### Participant 04

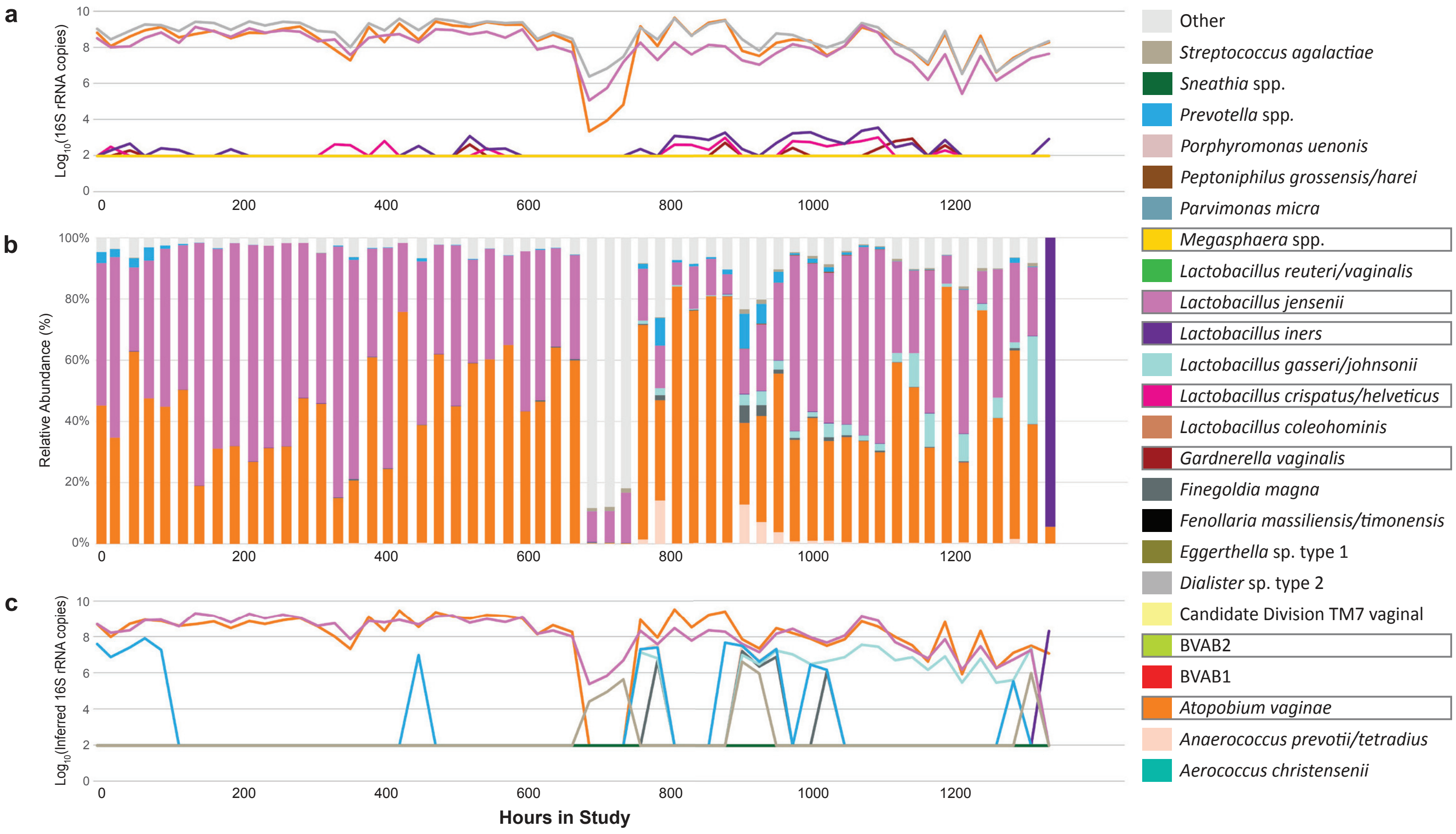

### Participant 05

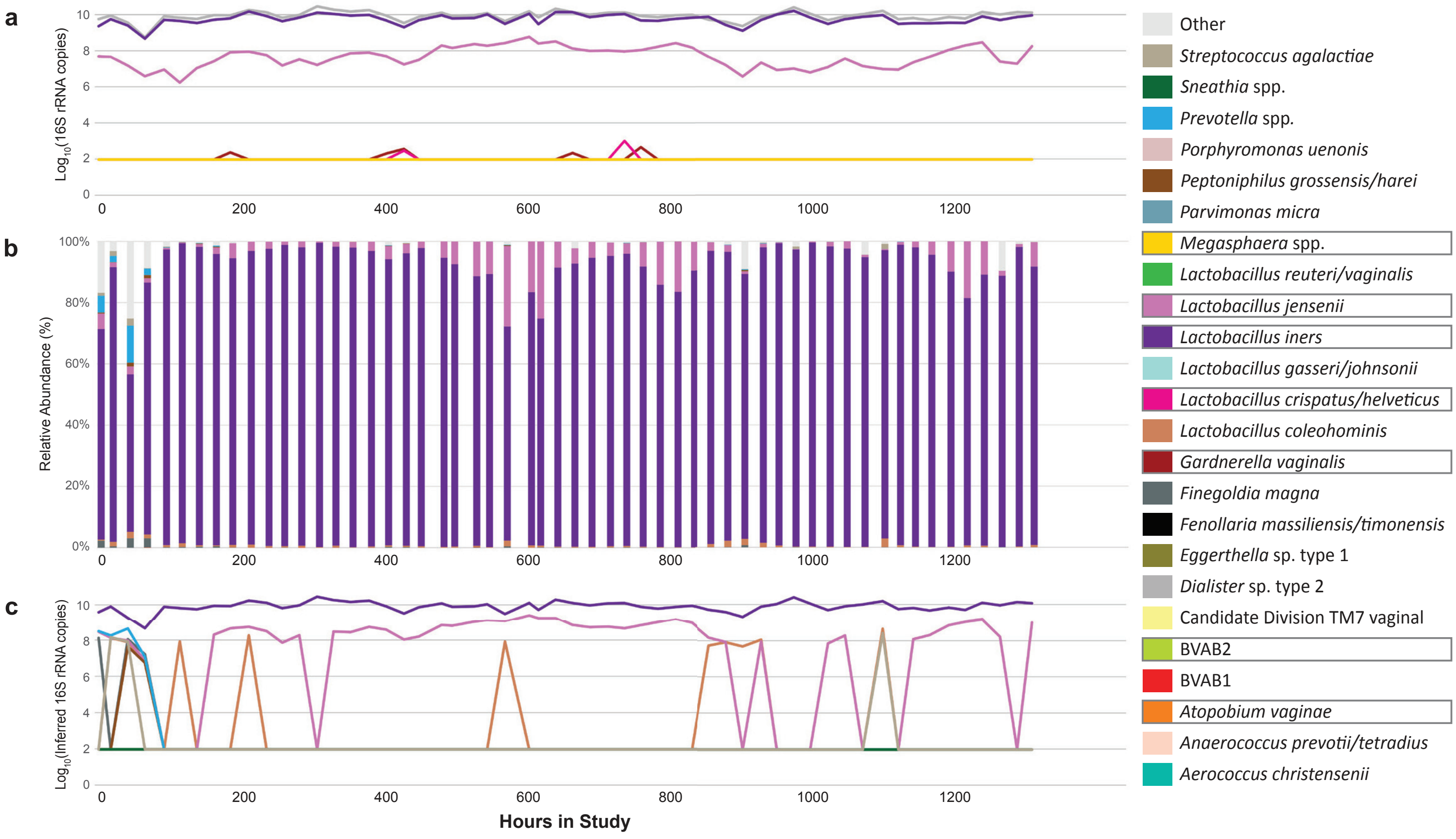


### Participant 06

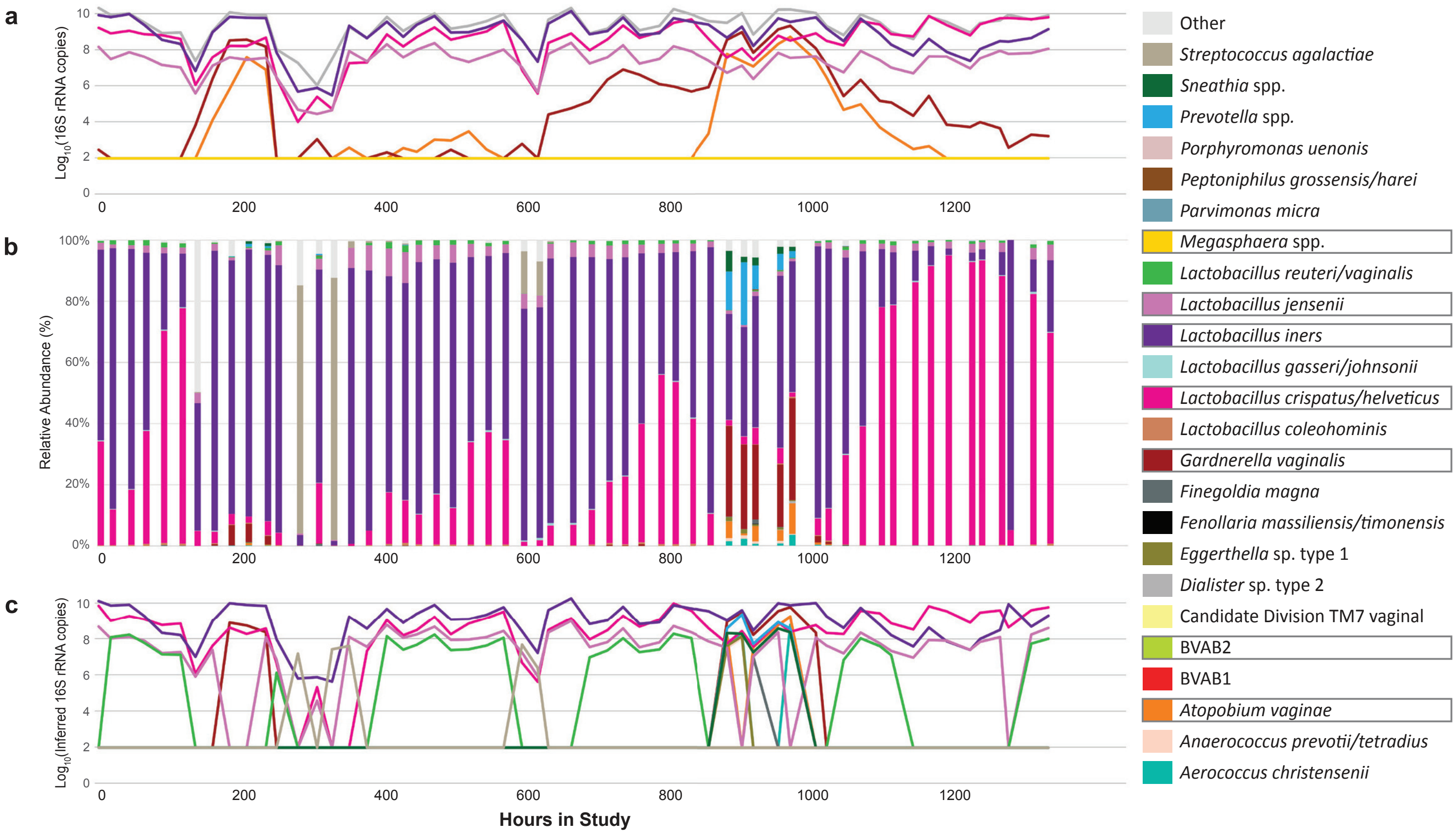

#### Participant 07

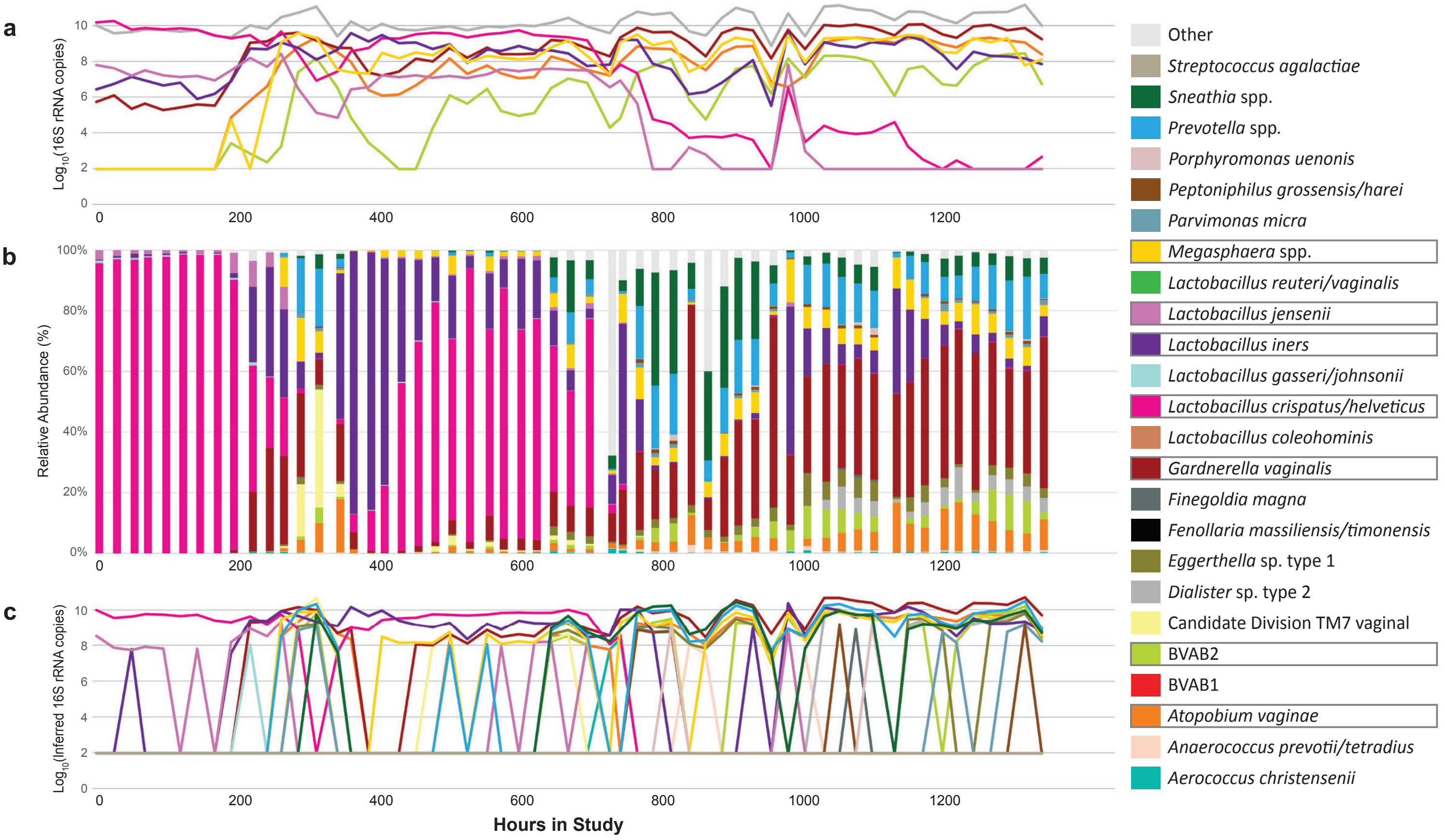


#### Participant 08

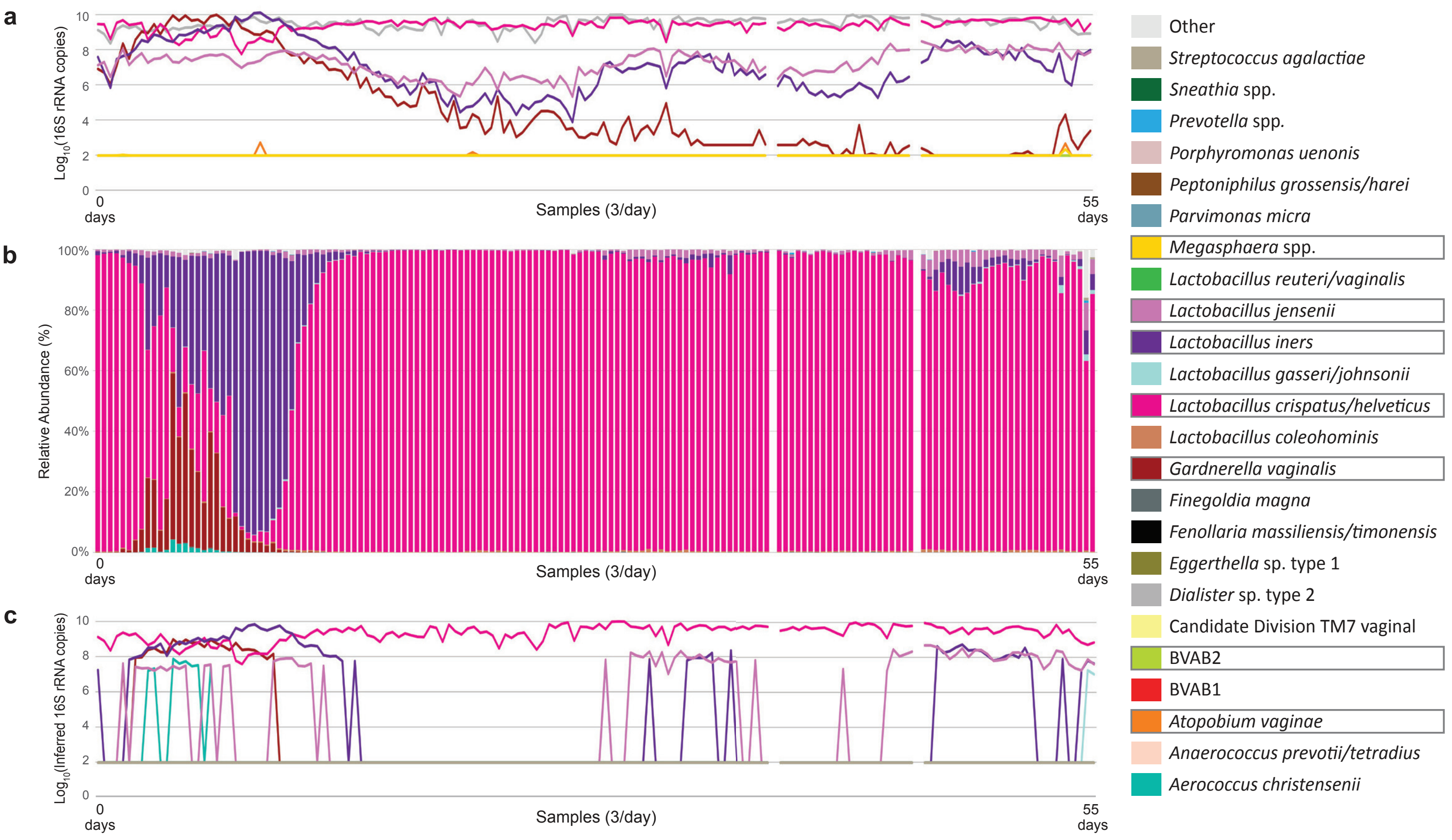

#### Participant 09

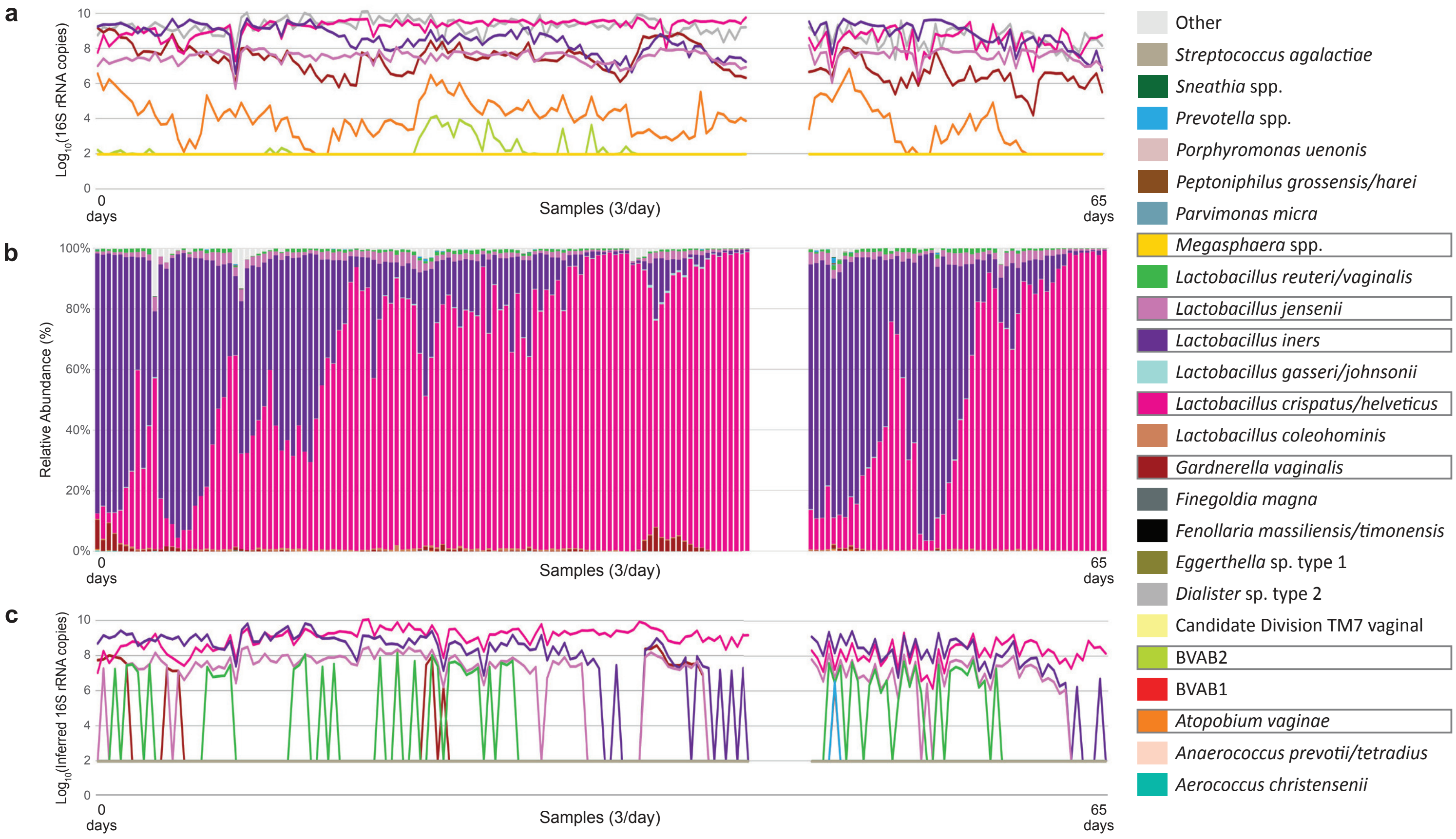


#### Participant 10

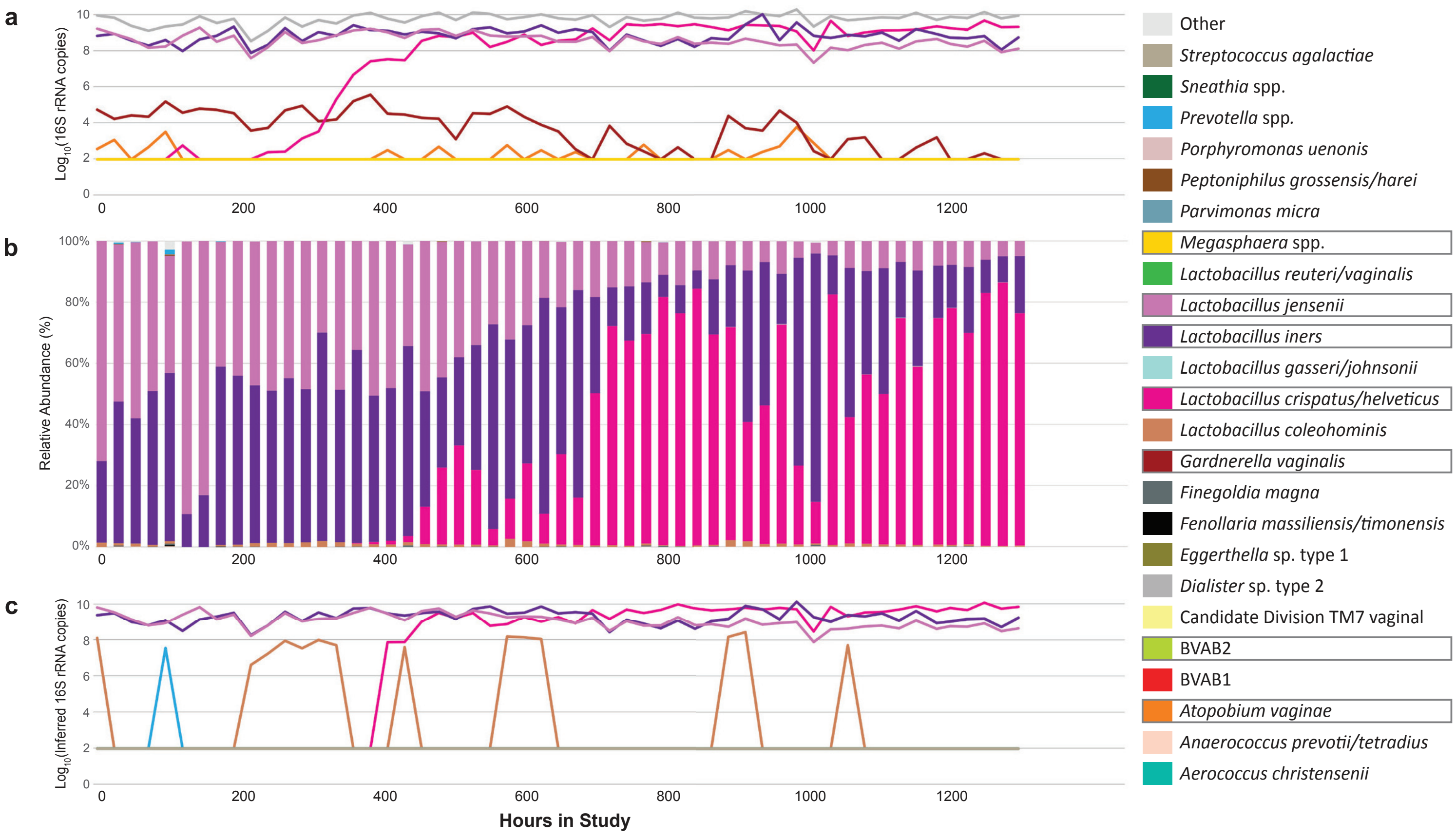

#### Participant 11

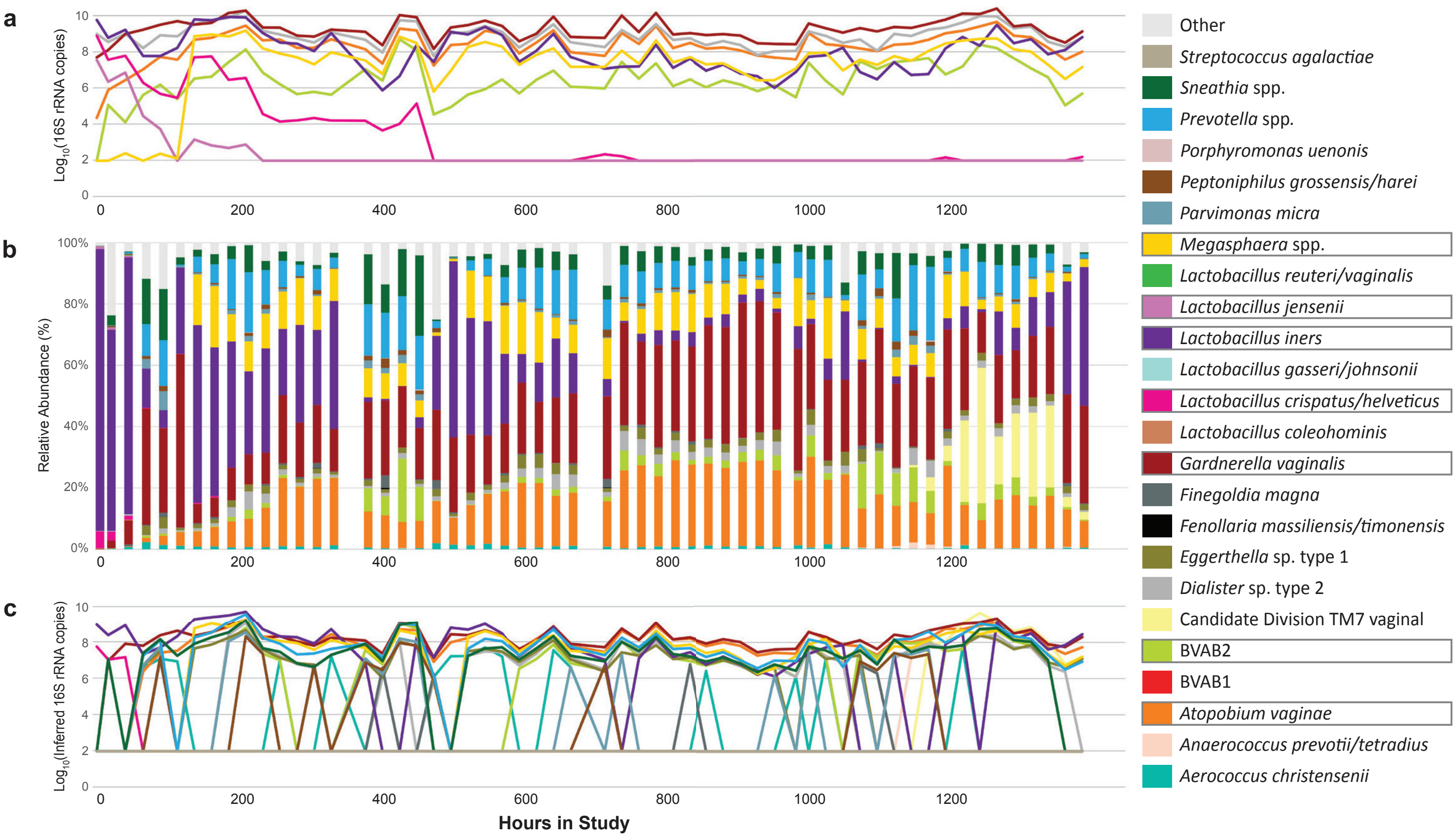


#### Participant 12

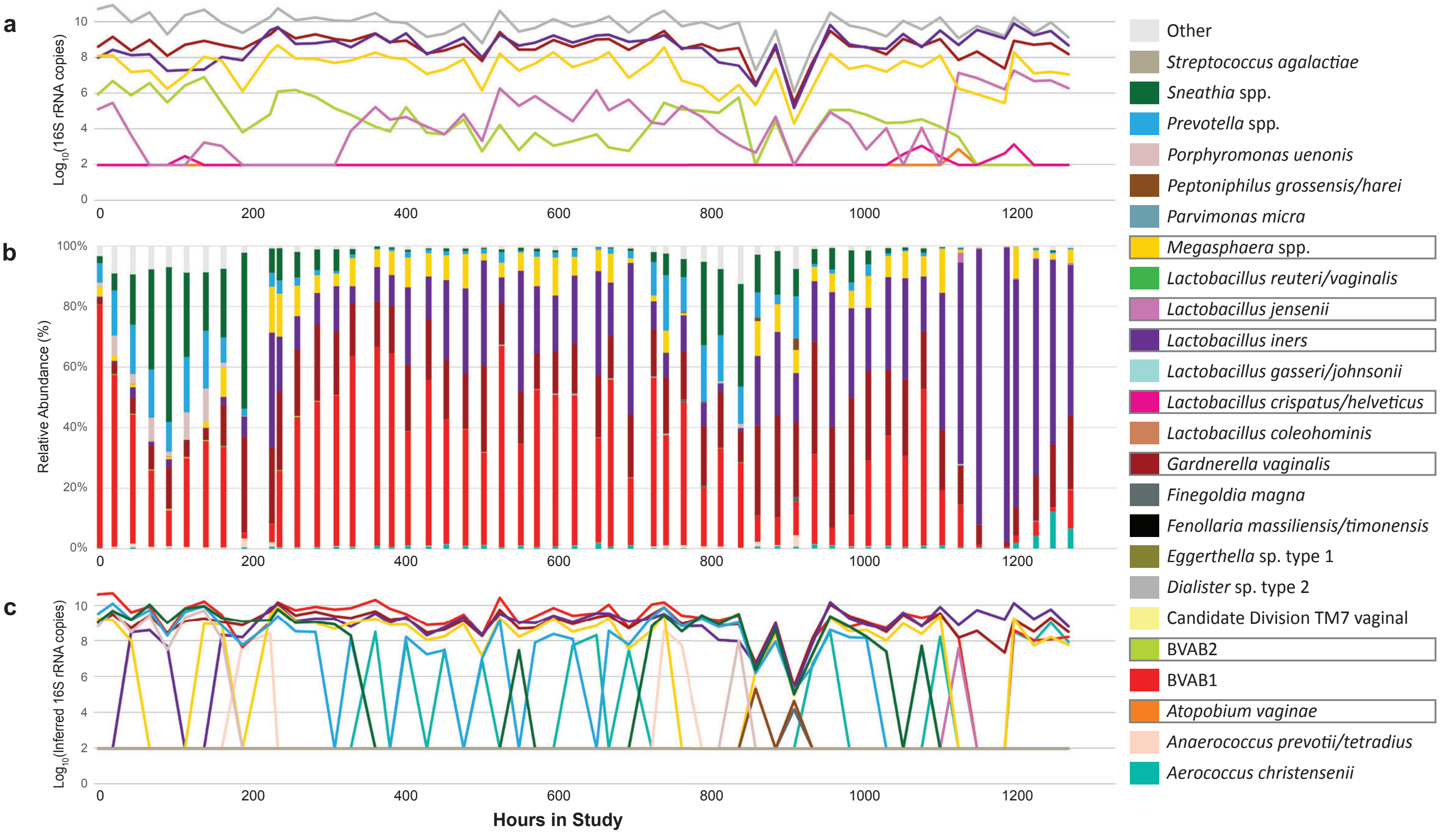

### Participant 13

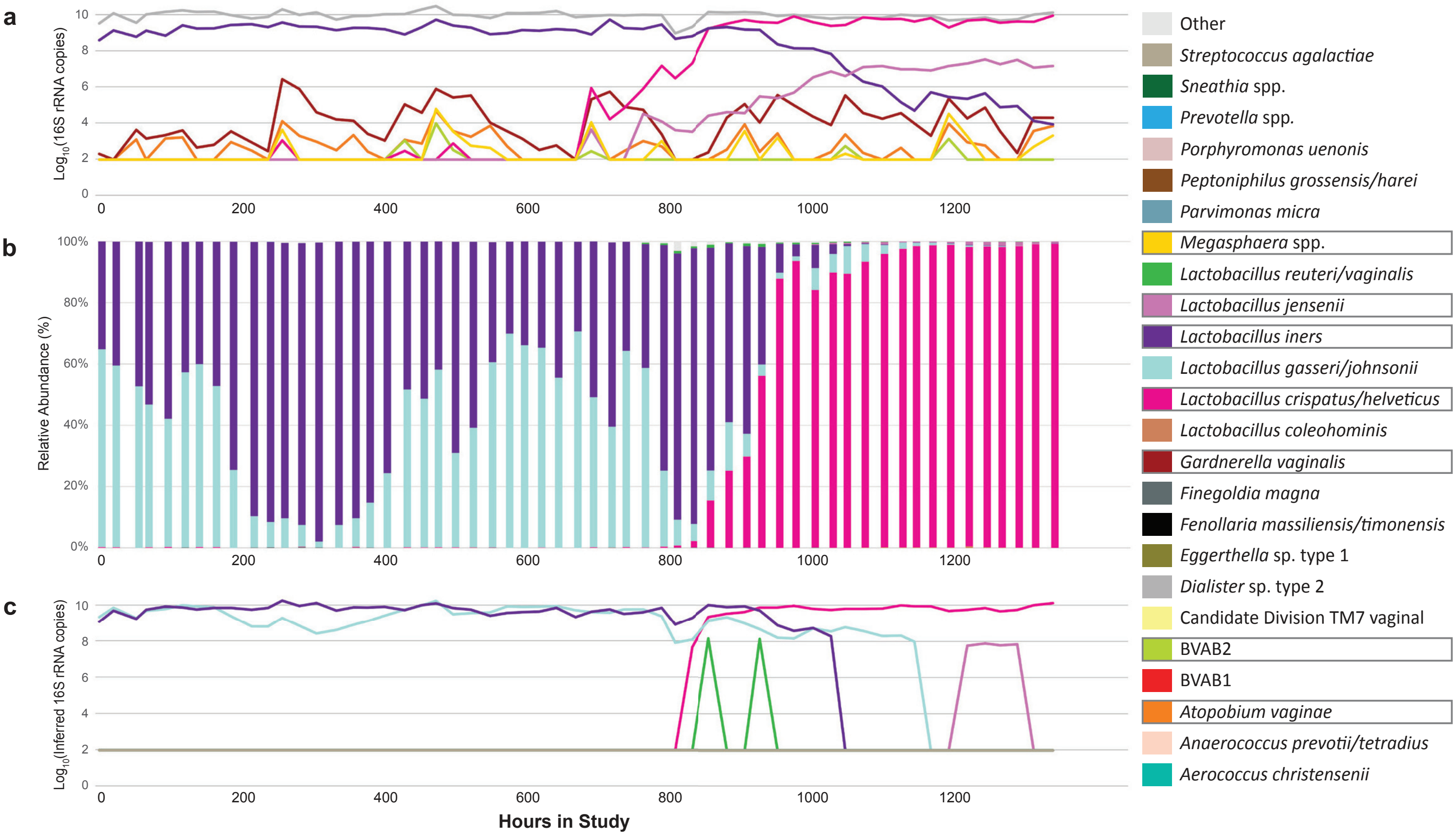


### Participant 14

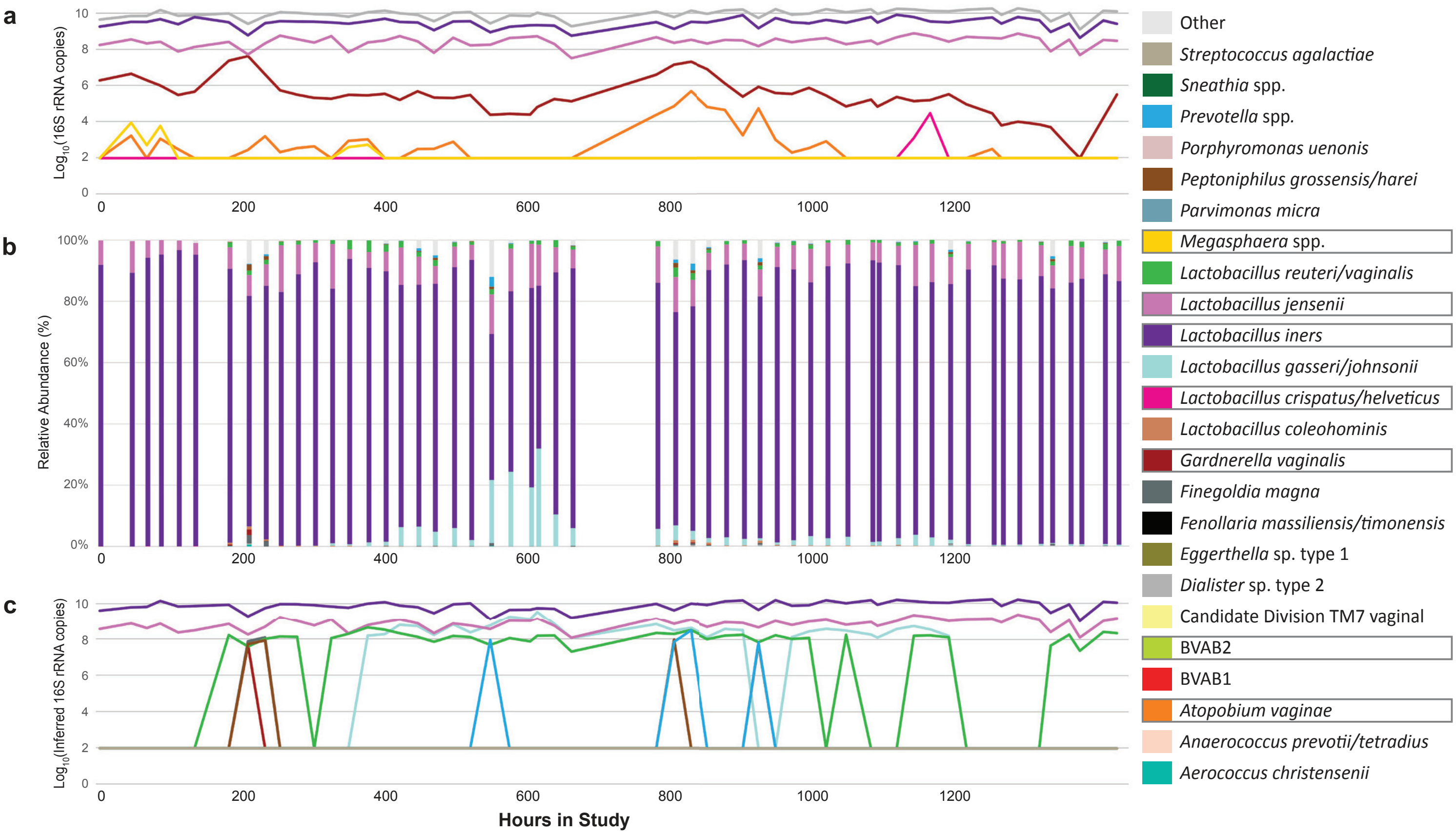

#### Participant 15

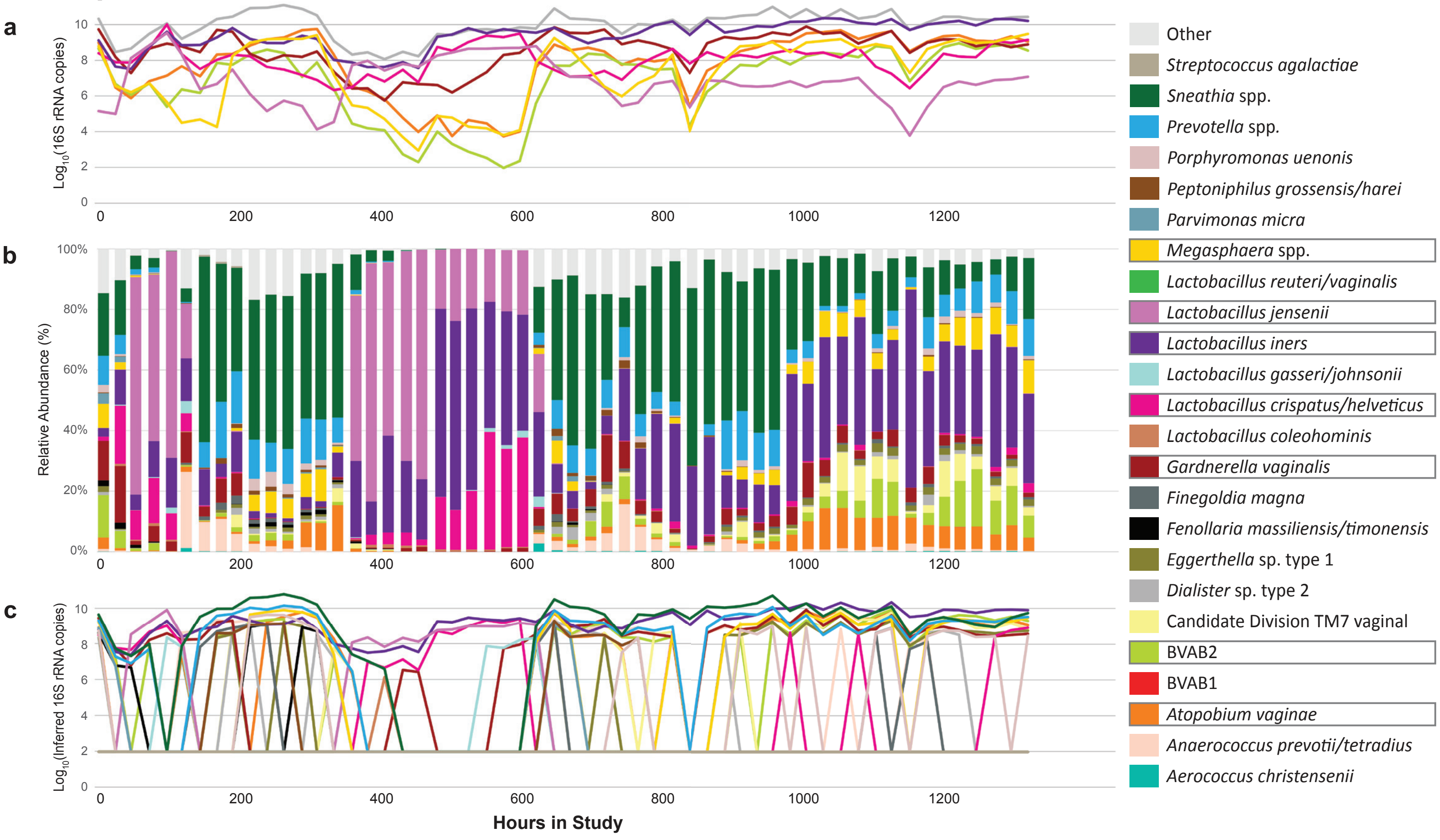


#### Participant 16

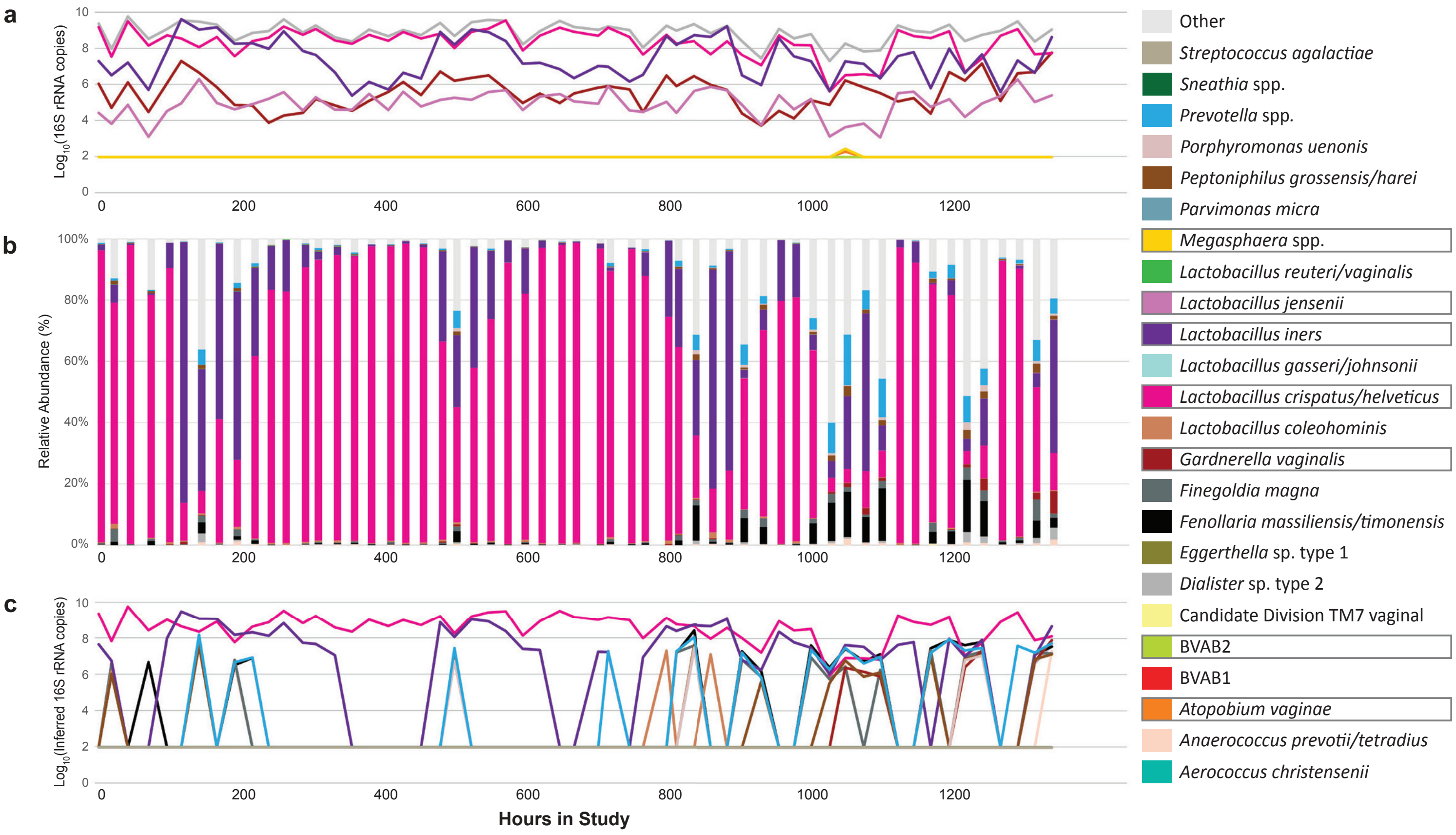

#### Participant 17

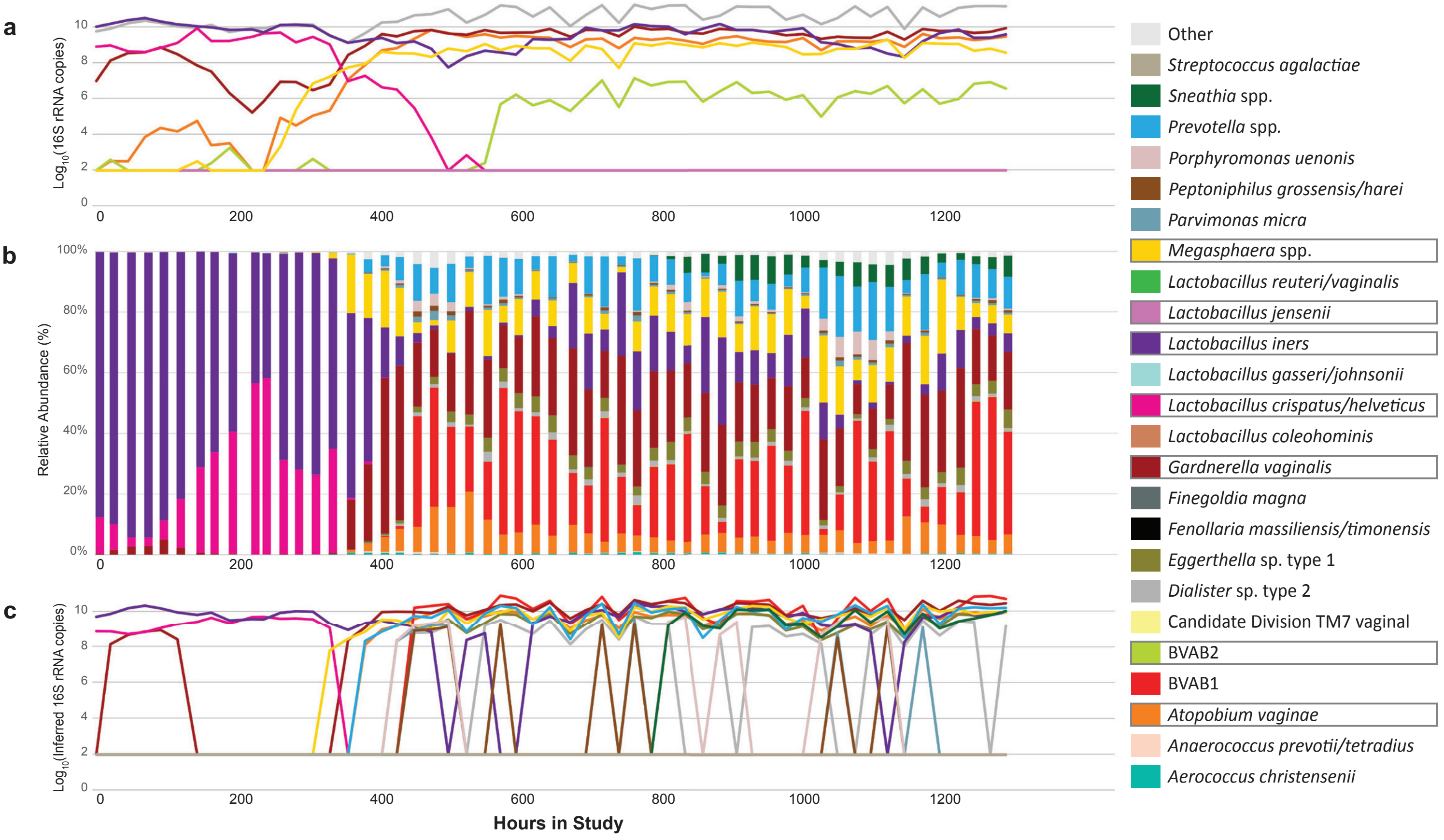


#### Participant 18

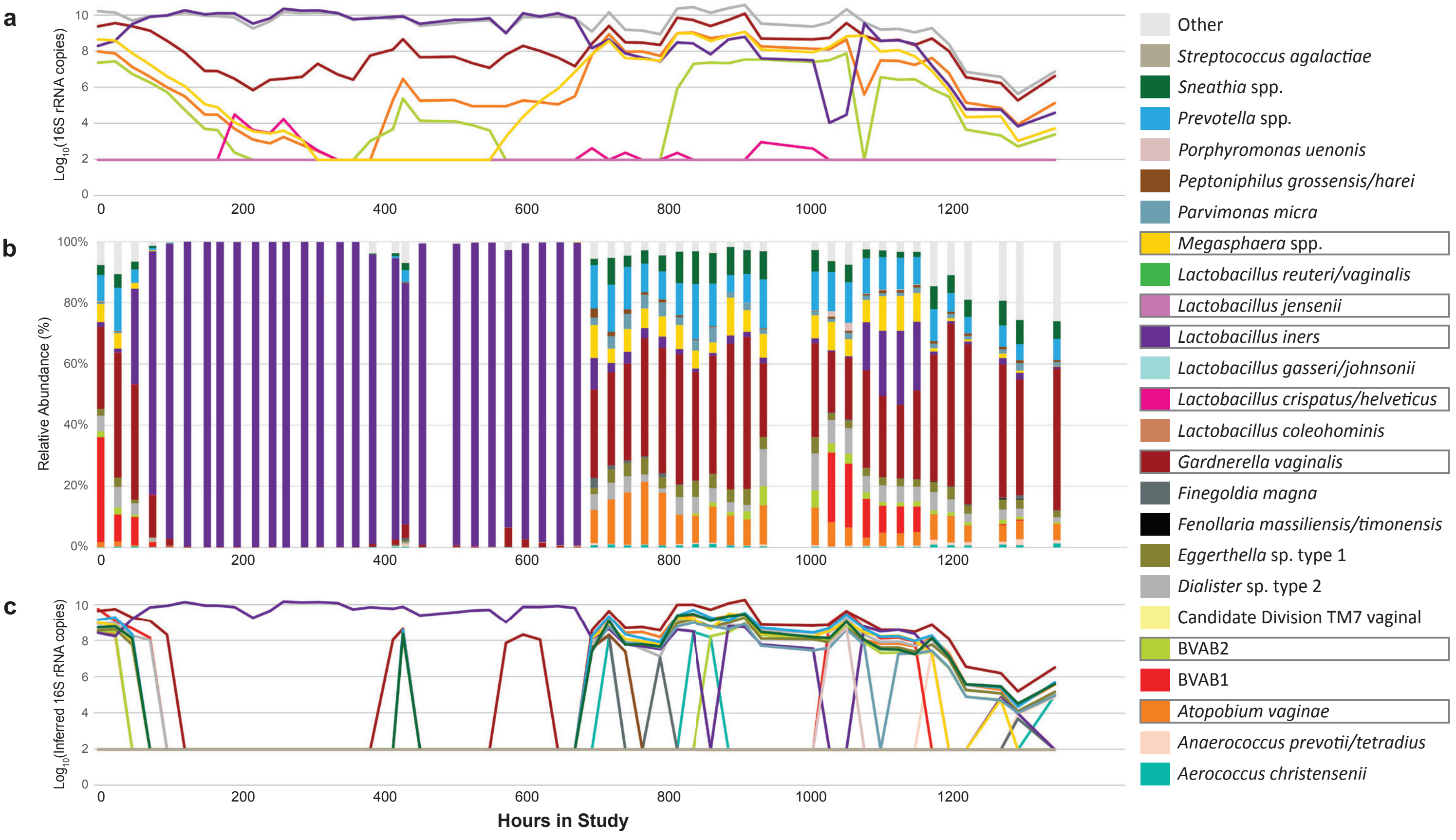

#### Participant 19

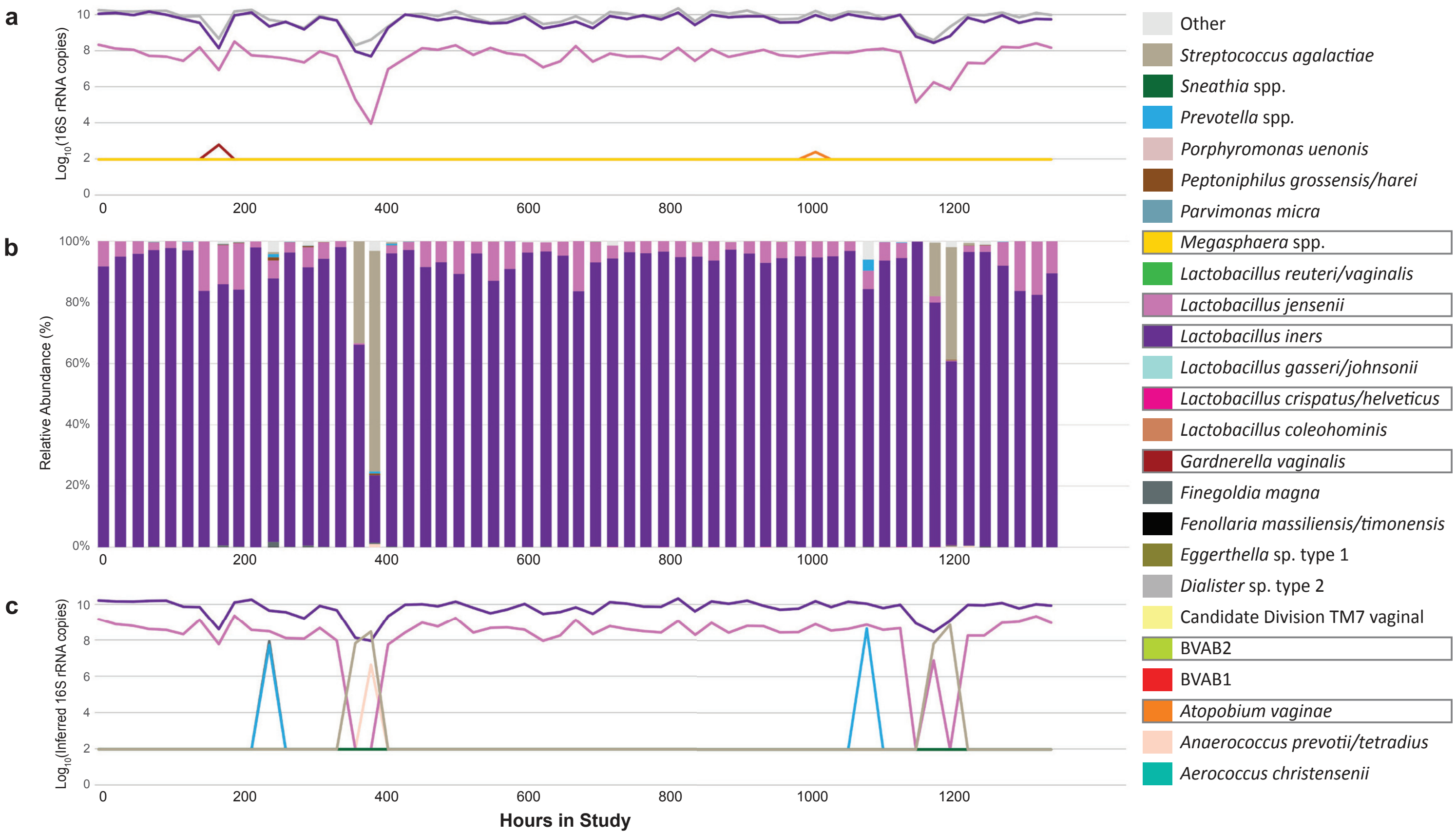


#### Participant 20

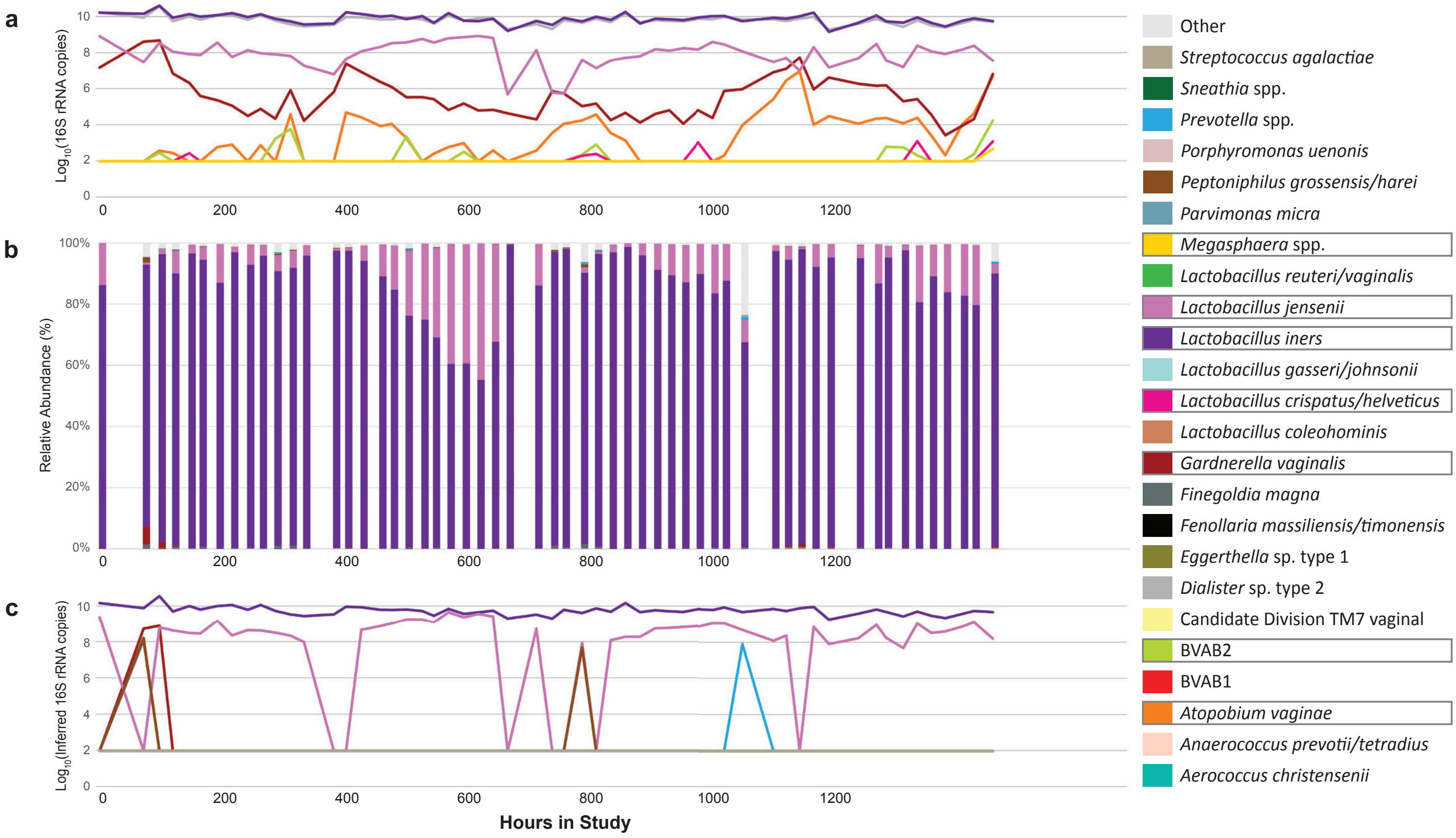

**Figure S2. Relative abundances estimates can misrepresent actual concentrations due to shifts in total bacterial load.** Vertical bars show relative abundance (%, left-y-axis), solid lines are absolute concentrations measured by qPCR and grey line is total bacterial load (both right y-axis). The dashed black line indicates detection threshold for qPCR data (93.8 16S rRNA copies). Arrows indicate obvious timepoints when relative abundance changes are discordant from absolute abundance changes, often when bacterial loads shift dramatically, or relative abundance is low. All seven species measured with qPCR in the study are included in separate panels from individual participants. a*)* *L. jensenii*, Participant 20 b) *L. iners*, Participant 02 c) *G. vaginalis*, Participant 18 d) BVAB2 Participant 15 and e) *A. vaginae* Participant 18.

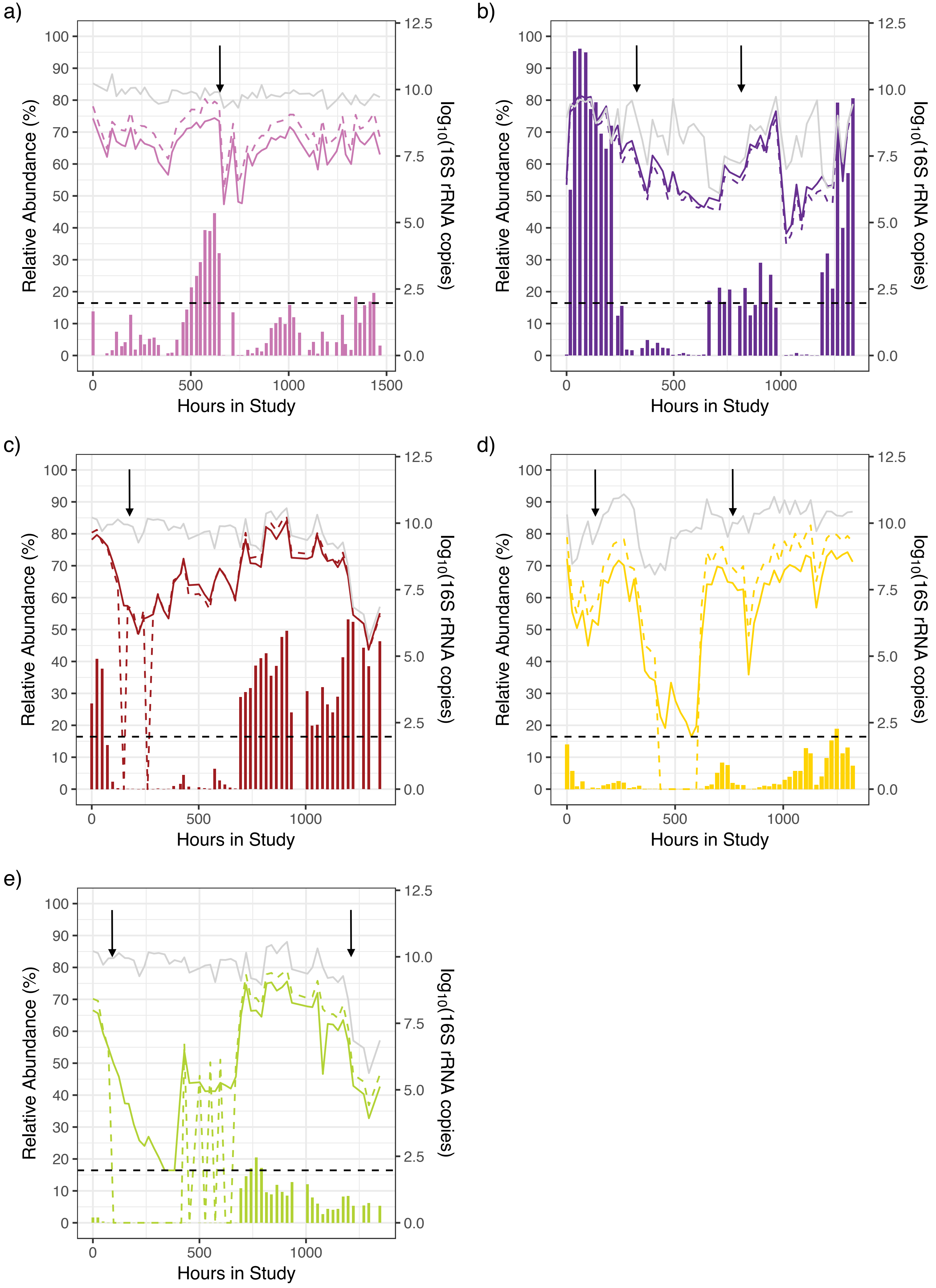
a)
b)
c)
d)
e)
Relative Abundance (%)
log10(16S rRNA copies)
Hours in Study
0
500
1000
1500
100
90
80
70
60
50
40
30
20
10
0.0
2.5
5.0
7.5
10.0
12.5

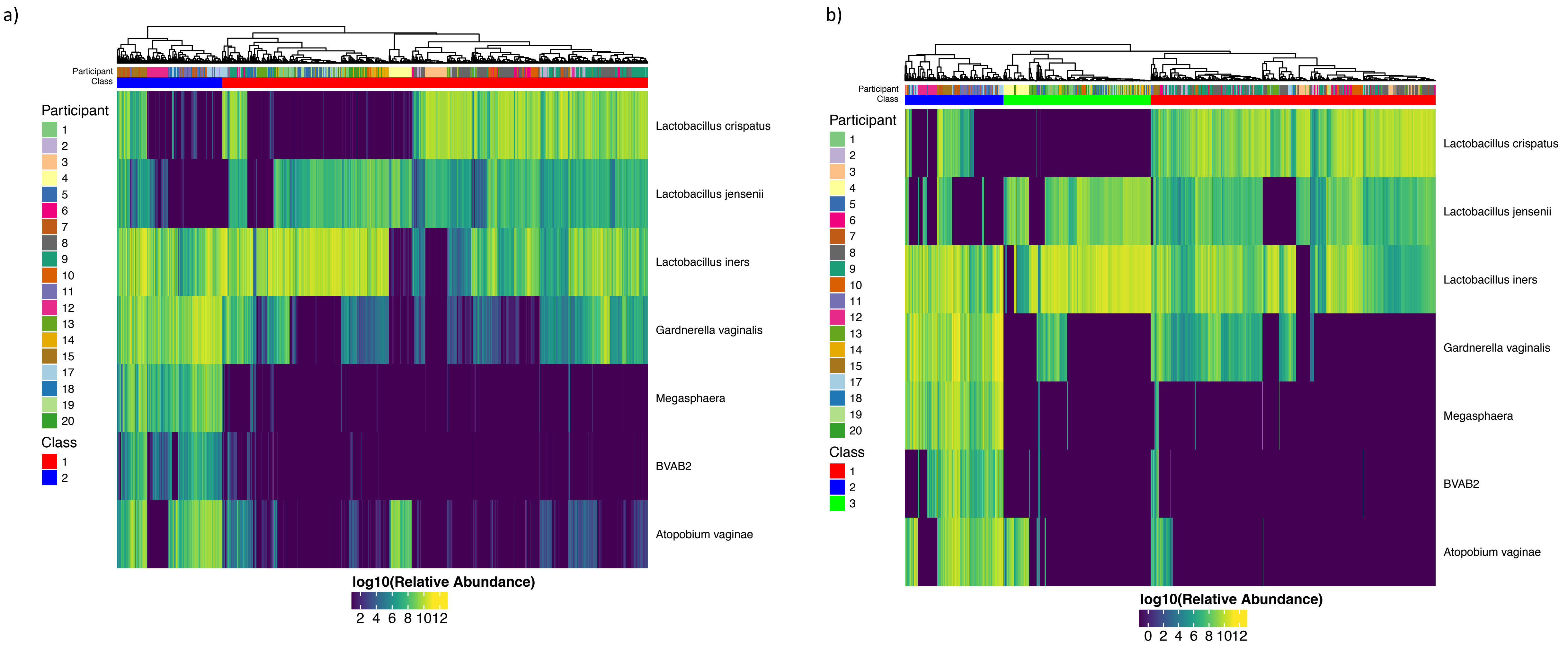


**Figure S3.** Complete hierarchical linkage clustering by 7 key species across all samples a) absolute concentration and b) inferred concentration. The two analysis are somewhat coherent with one another. Although the two dendrograms are in agreement with a low entanglement coefficient 0.11 the measurements identify a different number of clusters. The third cluster identified by inferred concentration, appears to be primarily distinguished from other samples dominated by lactobacilli species (Class 1), by having a low concentration of *L. crispatus* relative to other samples dominated by Lactobacilli species. This occurs as the inferred concentrations can report lower concentrations than the threshold of detection reported by absolute concentrations.

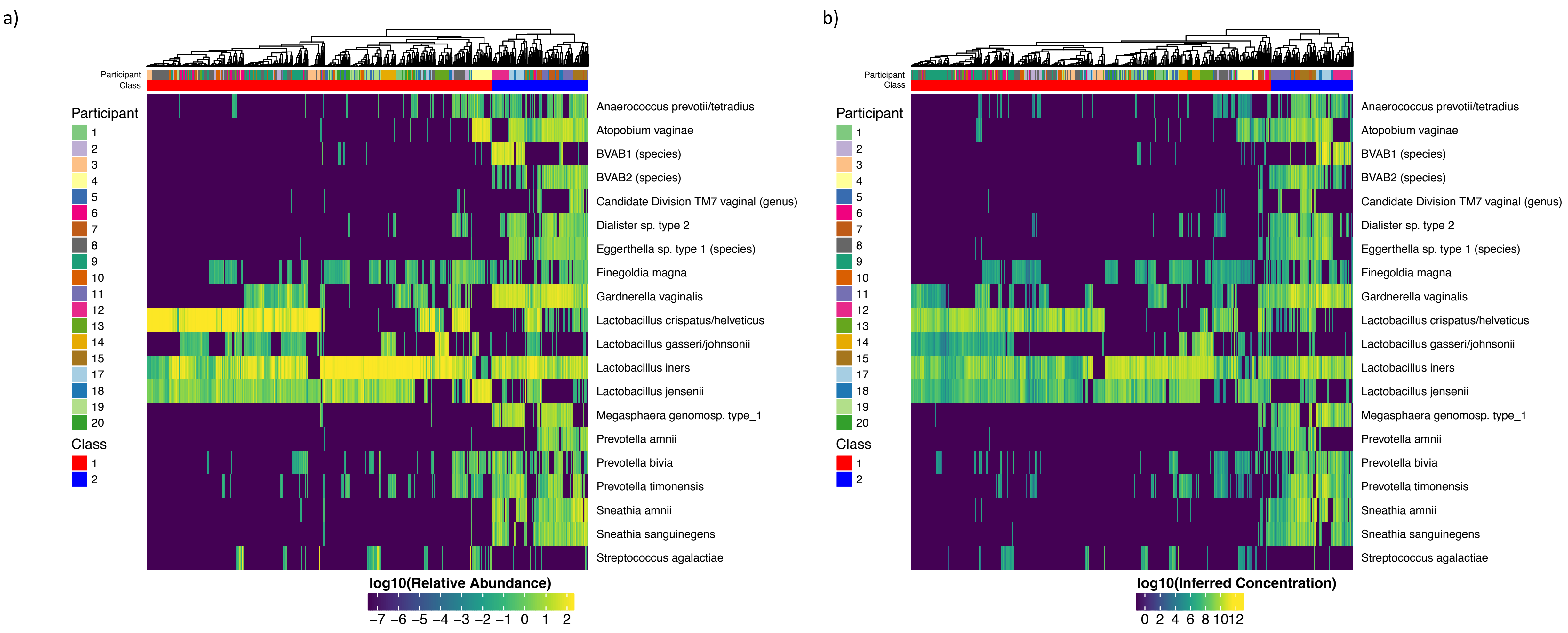


**Figure S4.** Complete hierarchical linkage clustering using the top 20 most abundant species across all samples a) relative abundance and b) inferred concentration. The two analysis are coherent with one another. Specifically, the two dendrograms are in agreement with a low entanglement coefficient 0.12 and both measurements identify the same number of clusters.
